# Comparative profiling of N-respirasomes predicts aberrant mitochondrial bioenergetics at single-cell resolution

**DOI:** 10.1101/2020.12.07.414730

**Authors:** Fabio Bertan, Lena Wischhof, Enzo Scifo, Mihaela Guranda, Joshua Jackson, Anaïs Marsal-Cots, Antonia Piazzesi, Miriam Stork, Michael Peitz, Jochen H. M. Prehn, Dan Ehninger, Pierluigi Nicotera, Daniele Bano

**Author notes:** Correspondence: Dr. Daniele Bano, Deutsches Zentrum für Neurodegenerative Erkrankungen (DZNE) Venusberg-Campus 1, Gebäude 99, 53127 Bonn (Germany), Prof. Pierluigi Nicotera, Deutsches Zentrum für Neurodegenerative Erkrankungen (DZNE), Venusberg-Campus 1, Gebäude 99, 53127 Bonn (Germany).

## Abstract

Mitochondria sustain the energy demand of the cell. The composition and functional state of the mitochondrial oxidative phosphorylation system are informative indicators of organelle homeostasis and bioenergetic capacity. Here we describe a highly sensitive and reproducible method for single-cell visualization and quantification of mitochondrial respiratory supercomplexes as a novel means of measuring mitochondrial respiratory chain integrity. We apply a proximity ligation assay (PLA) and perform comparative studies of mitochondrial CI, CIII and CIV-containing supercomplexes (or N-respirasomes) in fixed human and mouse brain tissues, tumorigenic cells, iPSCs and iPSC-derived NPCs and neurons. Our optimized approach enables a quantitative *in-situ* assessment of even subtle mitochondrial lesions associated with aberrant respiration. By combining quantitative proteomics with single cell imaging analysis, we also report the mechanistic contribution of the MICOS complex subunit CHCHD3 in regulating N-respirasomes. Overall, our PLA-based profiling of N-respirasomes establishes a sensitive and complementary technique for detecting cell-type specific mitochondrial perturbations in fixed materials.

## Introduction

Mitochondria are double-membrane organelles that supply ATP and essential building molecules for cell growth, maintenance and division. In differentiated cells under normal aerobic conditions, oxidative phosphorylation (OXPHOS) accounts for approximately 90% of the total required cellular energy (Wallace, 2018). Three complexes (i.e., CI, CIII and CIV) of the OXPHOS system couple the electron transfer to proton translocation across the inner mitochondrial membrane, thereby creating an electrochemical gradient that the ATP synthase (or complex V) utilizes for ATP production. Although individual respiratory complexes can randomly diffuse and transiently interact (Hackenbrock et al., 1986), they can also associate in stable, higher-order structures in equilibrium within the inner membrane of the cristae. It has been suggested that these “supercomplexes” (Schagger and Pfeiffer, 2000) may promote stability, kinetic advantages and functional flexibility of proton-pumping complexes (Acin-Perez et al., 2008; Calvo et al., 2020; Garcia-Poyatos et al., 2020; Lapuente-Brun et al., 2013; Letts et al., 2019; Wu et al., 2016). Although the physiological role remains a topic of intense scientific debate, it appears that superassembled complexes enhance respiratory capacity and limit the formation of byproducts, such as reactive oxygen species (ROS) (Calvo et al., 2020; Lapuente-Brun et al., 2013; Maranzana et al., 2013). It is less clear whether these higher-order respiratory supercomplexes (RSCs) enable a more efficient electron transfer and substrate channeling function (Bianchi et al., 2004; Calvo et al., 2020; Fedor and Hirst, 2018; Lapuente-Brun et al., 2013; Letts et al., 2019). In terms of organization, the NADH-respirasome (or N-respirasome) is an assembly as large as 19 nm in height and 30 nm in length, consisting of CI, CIII and CIV in a ratio of 1:2:1 (Calvo et al., 2020; Wu et al., 2016). The remaining fractions of CI and CIV can also organize in alternative assemblies with CIII, further expanding the array of possible quaternary species (Calvo et al., 2020; Cogliati et al., 2016; Lapuente-Brun et al., 2013; Letts et al., 2019). Based on recent evidence, CIII_2_ is central for both electron transport chain (ETC) homeostasis and RSC biogenesis, since it recruits components of nascent CI and CIV and acts as a primary seed that positively stimulates the full assembly and stability of individual complexes (Calvaruso et al., 2012; Protasoni et al., 2020). As originally postulated (Schagger and Pfeiffer, 2000), the presence and ratio of different RSCs depend on various factors (e.g., background, polymorphism, epigenetic regulation of gene expression) as well as tissue composition and bioenergetic demands of individual cells (Calvo et al., 2020; Cogliati et al., 2016; Lapuente-Brun et al., 2013; Sun et al., 2016). Functional defects, environmental toxins and genetic lesions that impair the expression and incorporation of ETC components can irremediably undermine the proper formation of respiratory complexes and, as a consequence, of functional respirasomes (Acin-Perez et al., 2008; Calvo et al., 2020; Cogliati et al., 2016; Lapuente-Brun et al., 2013; Protasoni et al., 2020). Since aberrant mitochondrial bioenergetics often lead to metabolic syndromes, inherited pathologies and debilitating neurodegenerative diseases (Area-Gomez and Schon, 2014; Bano and Prehn, 2018; Frazier et al., 2019; Gorman et al., 2016), the assessment of the RSC in patient-derived materials may provide additional informative cues about disease etiology.

In biomedicine, a range of cellular parameters have been used as proxy markers of mitochondrial function (Connolly et al., 2018; Frazier et al., 2020). In the last two decades, innovative approaches have been developed for population-based quantifications of mitochondrial bioenergetics, of which one prevailing functional read-out is the amount of O_2_ consumed over time. The most advanced systems measure oxygen consumption rate (OCR) using conventional O_2_ electrode chambers (e.g., Hansatech Oxygraph), O_2_-sensitive fluorescent indicators (e.g., Seahorse XF flux analyzer, Luxcel Bioscience’s MitoXpress) and amperometric O_2_ sensors (e.g., Oroboros Oxygraph-2k system) (Connolly et al., 2018). Despite the robustness and reliability of OCR measurements, these methods show several limitations in cell-specific assessment of mitochondrial performance in complex tissues such as the brain, unless laborious cell sorting precedes any analysis. To overcome this drawback, time-lapse fluorescent imaging analysis of various mitochondrial parameters (e.g., membrane potential, ROS, Ca^2+^signaling, *in situ* respiration) can effectively be performed in living tissues, as long as cell-type specific labelling (e.g., overexpression of a fluorescent protein as a marker) is available. Because none of these methods can be applied to fixed materials, one standard and widely accepted approach involves stainings of distinct ETC subunits in combination with markers that allow for discrimination of probe signals in a cell-type specific manner. The sequential analysis using selected and properly validated antibodies against subunits of the five OXPHOS complexes can enable the quantification of eventual changes in mitochondrial function. While this method is sufficiently accurate in the case of mitochondrial lesions associated with human pathologies (Lake et al., 2016; Wallace, 2018), it becomes less adequate to quantitatively measure adjustments due to biological processes that require the fine tuning of mitochondrial respiration, as in the case of cell differentiation or tumor formation. Similarly, this approach has limitations when OXPHOS assembly intermediates are still present in cells carrying lesions of unknown nature or upon exposure to toxins that interfere with mitochondrial biology. Other technical drawbacks include intrinsic experimental variability (i.e., instrument set-up and imaging acquisition parameters, sample collection and preparation) and eventual age-dependent accumulation of endogenous pigments (e.g., melanin) and lipochromes (e.g., lipofuscin) that may enhance autofluorescence background. Thus, a reliable *in situ* assessment of mitochondrial respiratory integrity in freshly prepared patient-derived samples or *postmortem* tissues still remains technically challenging, undermining diagnostic analyses as well as biomedical studies.

Starting with the knowledge that components of the OXPHOS system can associate in higher-order structures (Calvo et al., 2020; Cogliati et al., 2016; Lapuente-Brun et al., 2013; Schagger and Pfeiffer, 2000), we hypothesized that the abundance of RSCs might correlate with the integrity of the OXPHOS system and would sufficiently depict the state of mitochondrial respiration. Since the length of the mammalian N-respirasome (i.e., I_1_+III_2_+IV_1_) is approximately 30 nm (Wu et al., 2016) and, therefore, within the detectable range of a proximity ligation assay (PLA) (Fredriksson et al., 2002), we set out to develop an image-based protocol for measuring the abundance of CI,CIV-containing superassembled structures at single-cell resolution. We herein present our optimized, widely accessible and highly sensitive method that quantitatively profiles the state of the mitochondrial respiratory chain in fixed tissues.

## Results

### Proximity ligation-based staining allows the detection of mitochondrial superassembled complexes in fixed cells

We set off to design a reproducible and widely applicable proximity ligation-based method for visualizing and quantifying N-respirasomes at single-cell resolution (Figure 1A). We selected NADH dehydrogenase [ubiquinone] 1 beta subcomplex subunit 8 (NDUFB8) and cytochrome c oxidase subunit I (COX1 or MTCO1) because of their ubiquitous expression profiles across tissues (Calvo et al., 2016; Pagliarini et al., 2008). Primary antibodies against NDUFB8 and MTCO1 were chosen based on their high specificity and prior experimental validation in some of our works (Meyer et al., 2015; Troulinaki et al., 2018; Wischhof et al., 2018). As an initial cellular model, we employed near-haploid tumorigenic HAP1 cells. Confocal imaging analysis of NDUFB8 and MTCO1 revealed a clear co-localization with mitochondrial AIF and TOM20, respectively (Figure 1B-C). We validated our PLA conditions using different positive and negative controls (Figure 1D). In this regard, we observed signal when each primary antibody (i.e., NDUFB8 and MTCO1) was individually incubated with two oligonucleotide-labelled secondary antibodies that recognized the same IgG (Figure 1D, positive controls= +C). Conversely, no signal was observed in cells incubated with only one primary antibody and secondary antibodies against different IgGs or when the two secondary antibodies were incubated without primary antibodies (Figure 1D, negative controls= -C).

**Figure 1.**
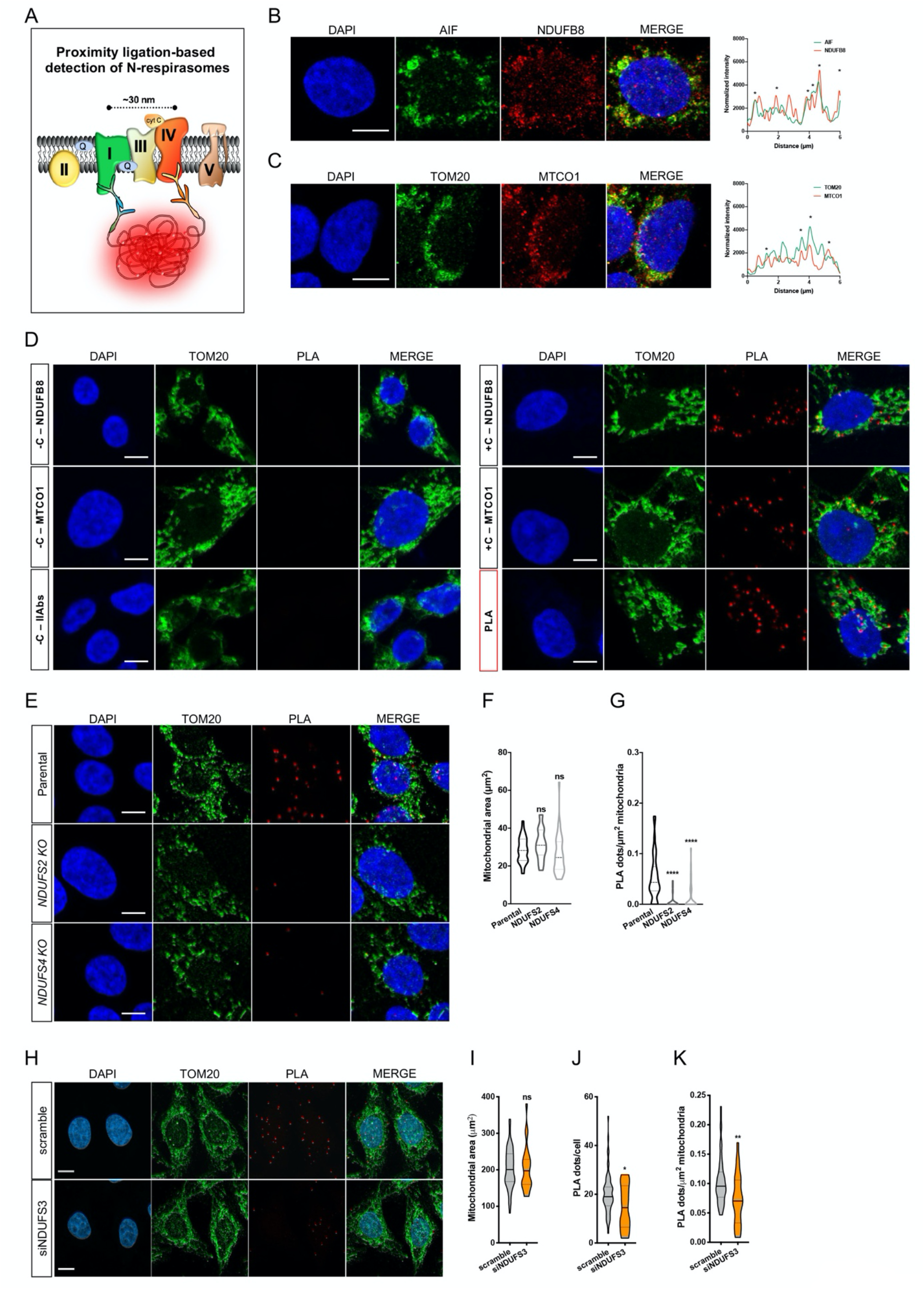
N-respirasomes are detectable in fixed cultured cells using PLA staining. (A) Scheme of the PLA-based approach to visualize N-respirasomes. Primary antibodies against nuclear encoded CI subunit NDUFB8 and mitochondrial DNA-encoded CIV subunit MTCO1 are used to detect superassembled structures. (B-C) Immunostaining and fluorescence profile of (B) AIF (green) and NDUFB8 (red) and (C) TOM20 (green) and MTCO1 (red) in parental HAP1 cells (scale bar: 5 μm). Colocalization analyses are reported in the right panels (stars show the overlapping fluorescent signals relative to distance). (D) PLA-based detection of N-respirasomes (red dots) along with staining of the mitochondrial marker TOM20 (green) and the nuclear marker DAPI (blue). Negative and positive experimental controls are reported for NDUFB8, MTCO1 and respective secondary IgGs in HAP1 cells. Scale bar: 5 μm. (E) Representative confocal images of parental, *NDUFS2* KO and *NDUFS4* KO HAP1 cells stained with DAPI (nucleus, blue), TOM20 (mitochondria, green) and PLA (N-respirasomes, red). Scale bar: 5 μm. (F-G) Quantification of (F) mitochondrial area and (G) N-respirasomes normalized to mitochondrial area (n=3, parental = 37 cells, *NDUFS2* KO = 22 cells, *NDUFS4* KO = 37 cells, one-way ANOVA, \*\*\*\**p*<0.0001). (H-K) Representative images and quantification of (I) mitochondrial area (μm^2^), (J) PLA dots/cell and (K) PLA dots/μm^2^mitochondria in HeLa cells transfected with scramble and siRNA against *NDUFS3* (n=1, scramble = 60 cells, siNDUFS3 = 24 cells, Student’s *t*-test, \*\**p*<0.01, \**p*<0.05). Scale bar: 10 μM.

When PLA was carried out using both anti-NDUFB8 and anti-MTCO1 antibodies together with their corresponding species-specific secondary antibodies, fluorescent positive PLA dot-like structures were easily detectable, clearly quantifiable and primarily co-localized within mitochondrial TOM20-labelled structures (Figure 1D). Having established the optimal experimental conditions for our assay, we tested whether our method could detect changes in mitochondrial RSCs by using cells carrying genetic lesions of the OXPHOS system. Since CI mutations inhibit the formation of N-respirasomes as previously described (Calvo et al., 2020), we quantified variations of N-respirasome content in *NDUFS2* and *NDUFS4* knockout (KO) HAP1 cells (Gioran et al., 2019). We found that, while TOM20-labelled mitochondrial network did not reveal obvious differences (Figure 1E-F), there were significantly fewer PLA dots in KO cells compared to parental controls (Figure 1E, 1G). To corroborate our findings in another cellular system, we transfected HeLa cells with scramble and small interfering RNA (siRNA) against *NDUFS3*. Compared to control, we found that transient downregulation of the CI subunit NDUFS3 did not alter mitochondrial area, whereas it decreased the number of PLA dots (Figure 1H-K). Together, this first set of validation data indicates that our proximity ligation-based staining is sufficiently sensitive to detect differences in N-respirasome levels in mitochondrial deficient cells.

### N-respirasome content diminishes in the brain of a mouse model of Leigh syndrome

We sought to evaluate the versatility of our PLA-based method in defining mitochondrial defects in neurons of brain tissues. To do so, we employed control (i.e., wild type: wt) and *Ndufs4* knockout mice (KO), the latter being a widely accepted preclinical model of Leigh syndrome (Kruse et al., 2008; Quintana et al., 2010). As previously shown, *Ndufs4* KO mice exhibit clear signatures of aberrant OXPHOS causally linked to the development of encephalomyopathy, ultimately resulting in premature death within 7-10 weeks of age (Kruse et al., 2008; Quintana et al., 2010). First, we measured OCR in *ex vivo* brain slices from 1-month-old control and *Ndufs4* KO mice, where we observed a tendency toward decreased basal respiration and response to mitochondrial inhibitors compared to controls (Figure 2A). Then, we separated proteins using conventional SDS-PAGE and performed immunoblot analyses. As expected, we observed a severe CI deficiency in samples from *Ndufs4* KO mice (Figure 2B). When we tested the loss of RSCs using BN-PAGE and subsequent immunoblot analysis of whole-brain lysates, we found a reduction of RSCs and a consequent increased amount of superassembled structures at lower molecular weights (Figure 2C). After validation and optimization of our antibodies in fixed brain tissues (Figure 2D-E), we ran high-resolution, Airyscan microscopy of double immunofluorescence staining, with PLA signals that co-localized with mitochondrial TOM20-labelled structures in neurons (Figure 2F). We carried out imaging analyses of hippocampal sections from age-matched *Ndufs4* KO mice and control littermates. PLA-based visualization of N-respirasomes was performed along with immunostaining of the neuronal marker NeuN and the mitochondrial marker TOM20 (Figure 2G-H). High-resolution confocal microscopy revealed that wt and *Ndufs4* KO CA1 neurons had a comparable somatic size (Figure 2I). However, KO cells exhibited a smaller mitochondrial network (Figure 2J) with a mitochondria-to-soma ratio that was statistically different to control neurons (Figure 2K). When normalized to the cell body or to mitochondrial area, the number of PLA dots was reduced by approximately half in *Ndufs4* KO CA1 neurons compared to control cells (Figure 2L-M). Linear correlation between PLA dots and mitochondrial area was observed only in control pyramidal cells, which was lost in *Ndufs4* KO CA1 neurons (Figure 2N). Together, our + findings indicate that *Ndufs4* KO neurons have a considerable loss of N-respirasomes compared to controls. Furthermore, our validated PLA-based method can quantify N-respirasome content at single-cell resolution in fixed brain sections.

**Figure 2.**
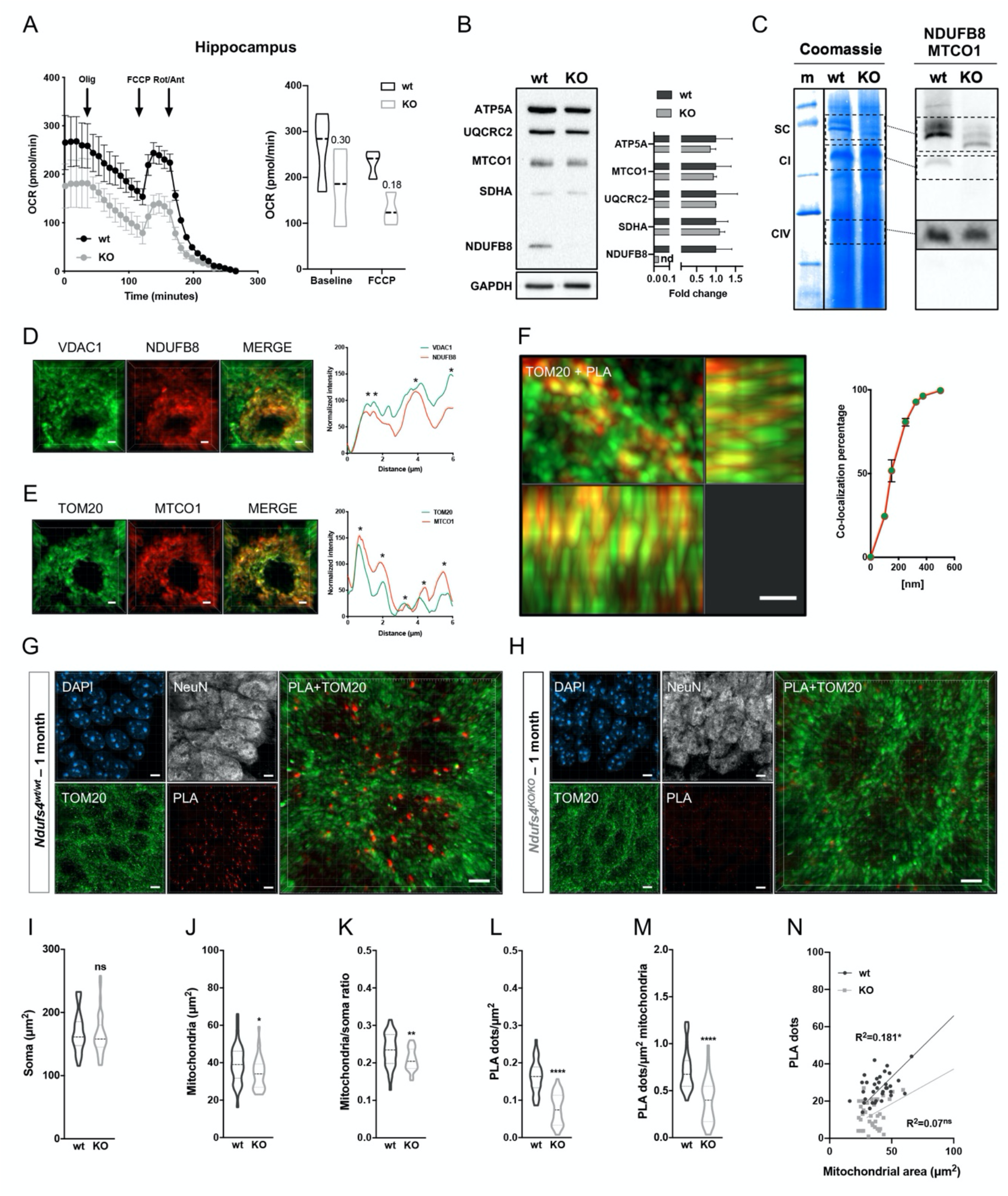
N-respirasome content diminishes in the brain of *Ndusf4* KO mice. (A) OCR measurement in *ex vivo* hippocampal brain sections from control (wt) and *Ndufs4* knockout mice (KO) (wt, n=3; *Ndufs4* KO, n=3, two-way RM ANOVA). (B) Immunoblot analysis of samples from wt and KO mice using antibodies against ETC subunits (CI, NDUFB8; CII, SDHA; CIII, UQCRC2; CIV, MTCO1; CV, ATP5A; wt, n=2; *Ndufs4* KO, n=2). GAPDH was used as loading control. Densitometry is relative to wt samples and reported as mean ± SEM (nd=not determined). (C) BN-PAGE and composite image of immunoblots obtained using antibodies against NDUFB8 (CI) and MTCO1 (CIV, cropped bands). Samples were from wt and KO brain homogenates (wt, n=3; *Ndufs4* KO, n=3). (D-E) Co-immunostaining and fluorescence profiles of (D) VDAC1 and NDUFB8 and (E) TOM20 and MTCO1 in control mouse brain sections (two-way ANOVA \**p*<0.05). Scale bar: 5 μm. (F) Co-localization between mitochondria (TOM20) and N-respirasomes (PLA). Scale bar: 1 μm. (G-H) Representative pictures of hippocampal CA1 pyramidal neurons from wt and KO mice stained with DAPI (nucleus), NeuN (neurons), TOM20 (mitochondria) and PLA (N-respirasomes). Scale bar: 5 μm. (I-N) Quantification of (I) area of the soma, (J) mitochondrial area, (K) mitochondria/soma ratio, (L) PLA dots normalized to the soma, (M) PLA quantification of N-respirasomes normalized to the mitochondrial area and (N) linear correlation of PLA dots and mitochondrial area (wt, n=3, 34 cells; *Ndufs4* KO, n=3, 37 cells). Data are represented as a violin plot with median [25^th^and 75^th^ percentile], Student’s t-test, n=3, \*\*\*\**p*<0.0001, \*\**p*<0.01, \**p*<0.05.

### Single-cell quantification of N-respirasomes enables comparative assessments of mitochondrial lesions during neurodegenerative processes

To perform comparative analysis of mitochondrial lesions in brains with distinct pathological profiles, we employed two additional mouse models of mitochondrial disease. Namely, we used Harlequin (Hq) mutant mice carrying a proviral insertion in the X-linked *Aifm1* gene (Benit et al., 2008; Klein et al., 2002; Wischhof et al., 2018) and *Aifm1 (R200 del*) knockin mice expressing a disease-associated variant of the apoptosis-inducing factor (AIF) (Wischhof et al., 2018). These two mouse models exhibit signatures of aberrant OXPHOS (Wischhof et al., 2018), however only Hq mutant mice display obvious gliosis and loss of cerebellar Purkinje cells at 6 months of age (Supplementary Figure S1A) (Klein et al., 2002; Wischhof et al., 2018). Because the Hq strain has a mixed genetic background (i.e., CBA/CaJ and C57BL/6J) (Klein et al., 2002), all analyses were strictly performed with the relative littermates. As a first step, we measured mitochondrial respiration in *ex vivo* brain sections (i.e., cerebella, hippocampi and cortices) using conventional Seahorse respirometry. Starting from comparable basal respiration levels, addition of mitochondrial respiratory inhibitors (i.e., oligomycin, FCCP and rotenone/antimycin A) resulted in similar responses, with only a minor tendency toward reduced OCR in AIF deficient tissues compared to controls (Supplementary Figure S1B-D). Using freshly prepared brain homogenates from Hq mutant, *Aifm1 (R200 del)* KI and control littermates, we performed SDS-PAGE and immunoblot analyses of OXPHOS components (Figure 3A and Supplementary Figure S1E-F). In line with our previous findings (Wischhof et al., 2018), we confirmed an evident loss of CI and CIII subunits in Hq mutant cerebella compared to controls, whereas only a moderate impairment of CI was observed in *Aifm1 (R200 del)* KI tissues (Figure 3A and Supplementary Figure S1E-F). In complex organs consisting of different cell types such as the brain, this set of data points out a technical limitation of OCR measurements in detecting mitochondrial defects associated with neurodegenerative processes.

**Figure 3.**
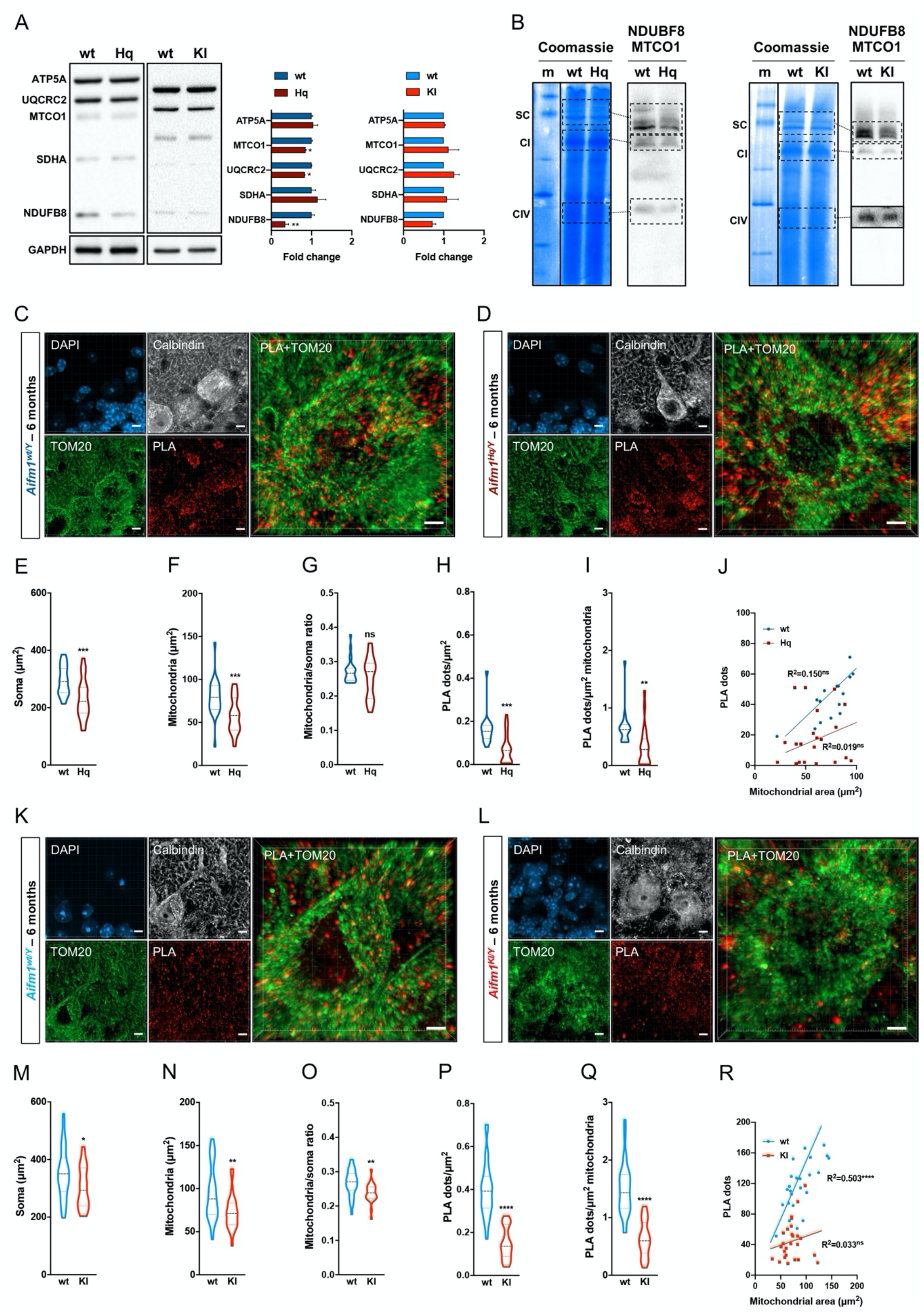
Single-cell quantification of N-respirasomes in Purkinje cells of Hq mutant and *Aifm1 (R200 del)* KI mice. (A) Immunoblot analysis of freshly frozen cerebella (wt, n=3; Hq, n=3; wt, n=3; KI, n=3) using antibodies against NDUFB8, SDHB, UQCRC2, MTCO1 and ATP5A. GAPDH was used as loading control. Densitometry is relative to wt samples and reported as mean ± SEM, Student’s t-test. (B) BN-PAGE and composite image of immunoblots using antibodies against NDUFB8 (CI) and MTCO1 (CIV, cropped bands for KI) using freshly frozen cerebella. (C-D) Representative pictures of cerebellar Purkinje cells from 6-month old control (wt) and Harlequin (Hq) mice stained with DAPI (nucleus), calbindin (Purkinje marker), TOM20 (mitochondria) and PLA (N-respirasomes). Scale bar: 5 μm. (E-J) Quantification of (E) area of the soma, (F) mitochondrial area, (G) mitochondria/soma ratio, (H) PLA-based dots normalized to the soma, (I) PLA-based dots normalized to the mitochondrial area and (J) linear correlation of PLA dots and mitochondrial area (wt, n=3, 16 cells; Hq, n=3, 22 cells). (K-L) Representative pictures of cerebellar Purkinje cells in control (wt) and *Aifm1 (R200 del*) KI mice stained with DAPI (nucleus), calbindin (Purkinje marker), TOM20 (mitochondria) and PLA (N-respirasomes). Scale bar: 5 μm. (M-R) Quantification of (M) area of the soma, (N) mitochondrial area, (O) mitochondria/soma ratio, (P) PLA-based dots normalized to the soma, (Q) PLA-based dots normalized to the mitochondrial area, (R) linear correlation of PLA dots and mitochondrial area (wt, n=3, 29 cells; KI, n=3, 26 cells). Data are represented as a violin plot with median [25^th^ and 75^th^ percentile], Student’s t-test, \*\*\*\**p*<0.0001, \*\*\**p*<0.001, \*\**p*<0.01, \**p*<0.05.

To ensure that AIF deficiency affects RSCs, brain homogenates were loaded on N-PAGE, blue Coomassie stained and then immunoblotted against anti-CI and anti-CIV antibodies (Figure 3B and Supplementary Figure S1G-H). We ascertained a consistent reduction of RSC content in AIF deficient cerebella relative to controls (Figure 3B). After a thorough validation of our antibodies and PLA protocol in cerebella slices (Supplementary Figure S2A-C), we went on to quantify N-respirasomes at single cell level. We applied our PLA protocol on cerebellar slices from 6-month-old Hq mutant, *Aifm1 (R200 del)* KI and relative littermates, then performed high-resolution confocal microscopy on calbindin-positive, TOM20-positive Purkinje neurons (Figure 3C-D and 3K-L). As a correlative observation, in Hq mutant cerebella we noticed an evident PLA signal from cells surrounding Purkinje neurons (Figure 3C-D), which is probably due to the large number of microglial cells in the degenerating tissue as previously reported (Klein et al., 2002; Wischhof et al., 2018). We performed a detailed quantification of mitochondrial area and N-respirasomes in calbindin-positive cells (Figure 3C-D and 3K-L). Compared to cells from control tissues, AIF deficient neurons had reduced soma size and mitochondrial area (Figure 3E-F, 3M-N), possibly more pronounced in KI cells since the mitochondria-to-soma ratio was statistically different to controls (Figure 3G and 3O). Importantly, PLA-based quantification (normalized to the area of the soma) revealed that N-respirasomes were diminished by 57% (Figure 3H) and 62.5% (Figure 3P) in Hq and *Aifm1 (R200 del)* KI Purkinje neurons, respectively, compared to controls. When PLA-dots were normalized to the mitochondrial area, N-respirasome content was decreased by 46.9% in Hq cells (Figure 3I) and by 57.8% in *Aifm1 (R200 del)* KI cells (Figure 3Q) compared to control neurons. A linear correlation between PLA dots and the mitochondrial area was observed only in tissues from C57BL/6NTac mice (Figure 3J and 3R), further emphasizing the contribution of the genetic background to RSC content as previously described (Calvo et al., 2020; Cogliati et al., 2016). Despite this intrinsic variability, our statistical analysis indicated that *Aifm1 (R200 del)* KI and, even more, Hq Purkinje cells display higher heterogeneity in term of N-respirasomes, with a subpopulation of neurons that have almost no detectable superassembled structures. In summary, our findings indicate that *in situ* quantification of N-respirasomes is an informative and sensitive approach for comparative analyses of mitochondrial defects in tissues undergoing neurodegenerative processes.

### The abundance of N-respirasomes decreases during aging

Having established the robustness of our method in mitochondrial mutant cells and tissues, we sought to perform experiments addressing open questions of biological interest. In this regard, we investigated the abundance of N-respirasomes in young versus old mice using our newly established approach. We employed horseradish peroxidase (HRP)-conjugated secondary antibodies in combination with a highly sensitive chromogenic substrate for bright-field visualization. As positive controls, each primary antibody was individually incubated with two oligonucleotide-labelled secondary antibodies (namely plus and minus) that recognized the same IgG. Following rolling circle amplification, antibody hybridization and enzymatic peroxidation, we imaged stained brain sections and detected strong and uniform dark signals as expected (Figure 4A). Conversely, no precipitate was observed in brain slices incubated without secondary antibodies (negative controls: Figure 4A). When PLA was carried out using NDUFB8 or MTCO1 and their corresponding species-specific secondary antibodies, we found that bright-field microscopy revealed more heterogeneous signals and intensity patterns in anatomically defined brain regions. Throughout the brain, PLA-positive dots were particularly evident in the cerebellar granule cell layer, in the stratum pyramidale of the hippocampus (i.e., DG, CA3 and CA1) and in the distal visual cerebral cortex (Figure 4A). We quantified the PLA signal in the cortex, cerebellum and hippocampus from 3-, 6- and 18-month-old C57BL/6J mice. Across the different areas, it appears that N-respirasomes significantly diminished between 3 and 6 months of age, with a more pronounced and consistent loss in older animals (i.e., 18-months of age) (Figure 4B-C). The age-dependent decrease of N-respirasomes was associated with a similar reduction of other mitochondrial superassembled species, as revealed by our BN-PAGE (Figure 4D). As a first line of evidence, we also performed PLA staining in hippocampal slices from a *postmortem* brain of a 71-year-old healthy donor (Supplementary Figure S3A-C). Both bright-field and fluorescent PLA staining of N-respirasomes were feasible in human derived samples. Together, this set of data indicates that N-respirasomes can be visualized in fixed brain tissues from aged organisms.

**Figure 4.**
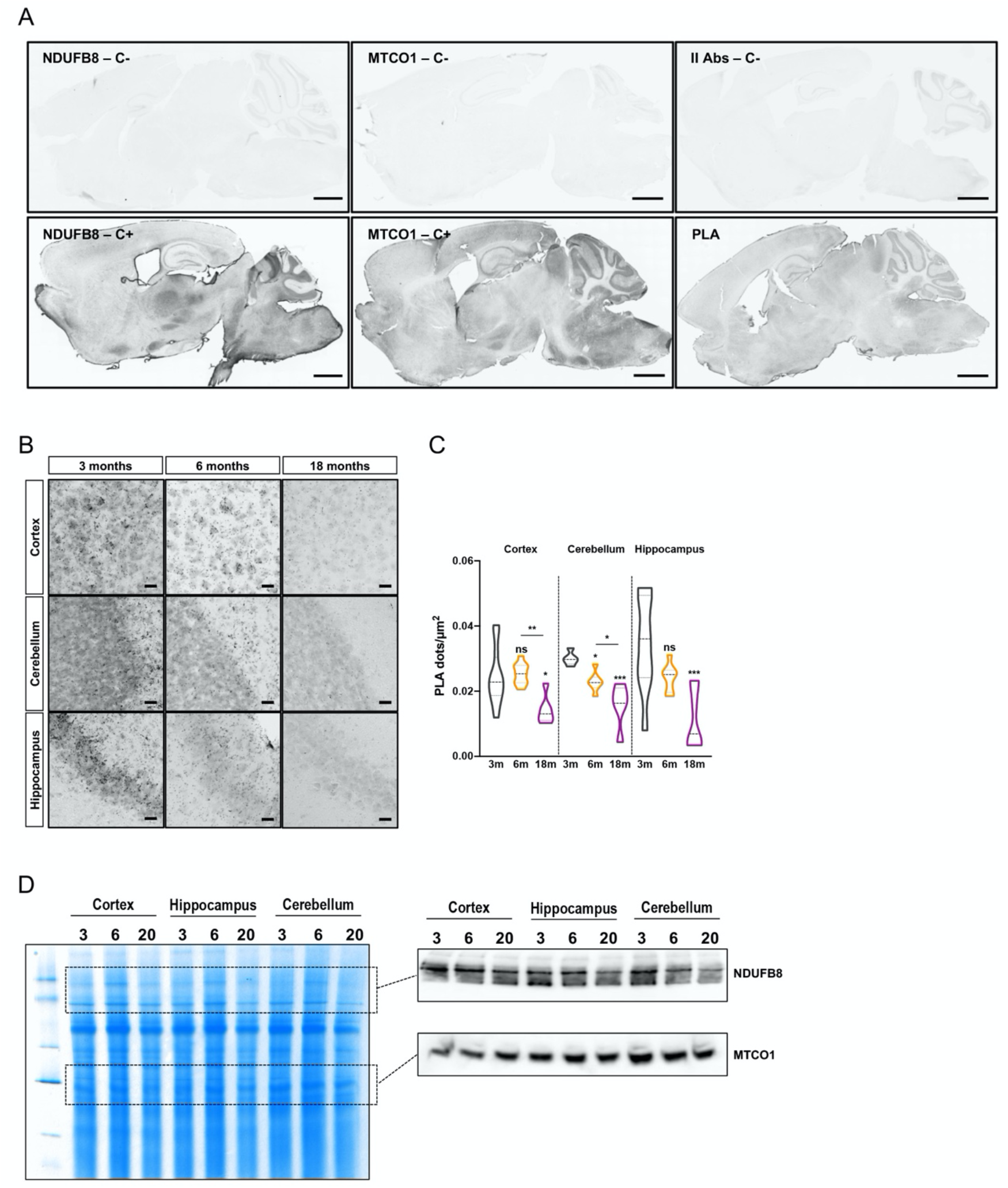
N-respirasome content decreases during aging in mouse brains. (A) Brightfield PLA-based detection of N-respirasomes in sagittal mouse brain sections. Negative and positive controls are for NDUFB8, MTCO1 with the corresponding combination of secondary IgGs. Scale bar: 1 μm. (B) High-resolution brightfield PLA staining of different brain areas. Scale bar: 50 μm. (C) Quantification of PLA dots/μm^2^ in cortexes, cerebella and hippocampi from 3-, 6-, 18-month old wt mice (n= 3). Data are represented as a violin plot with median [25^th^ and 75^th^ percentile], Student’s *t* test, \*\*\**p*<0.001, \*\**p*<0.01, \**p*<0.05. (D) BN-PAGE relative immunoblots (antibodies against NDUFB8, MTCO1) using freshly frozen mouse samples.

### Enhanced RSC content is associated with stem cell differentiation and survival

Since mitochondrial bioenergetics and its rewiring significantly influence stemness, cell lineage commitment and differentiation (Katajisto et al., 2015; Lisowski et al., 2018), we decided to compare RSCs in induced pluripotent stem cells (iPSCs) and iPSC-derived small molecule neural precursor cells (smNPCs). We profiled their OCR upon sequential addition of mitochondrial respiratory inhibitors (Figure 5A) and found that smNPCs had a much higher spare respiratory capacity, though reduced basal OCR, compared to iPSCs (Figure 5B-C). These differences in mitochondrial respiration were associated with variations in the composition of the OXPHOS system, including a two-fold increase of the CIII subunit UQCRC2 and a slight decreased expression of the CIV subunit MTCO1 (Figure 5D). BN-PAGE and subsequent immunoblots revealed a much higher content of RSCs in smNPCs compared to iPSC (Figure 5E). We applied our PLA-based method and found that N-respirasomes were much more abundant in smNPCs compared to iPSCs (Figure 5F-G), although their number substantially varied in individual cells, possibly because of the mixed population of cells at different cell cycle stages. Together, this line of evidence indicates that loss of stemness and consequent cell lineage commitment are linked to an evident remodeling of the OXPHOS system, with an enhanced formation of N-respirasomes.

**Figure 5.**
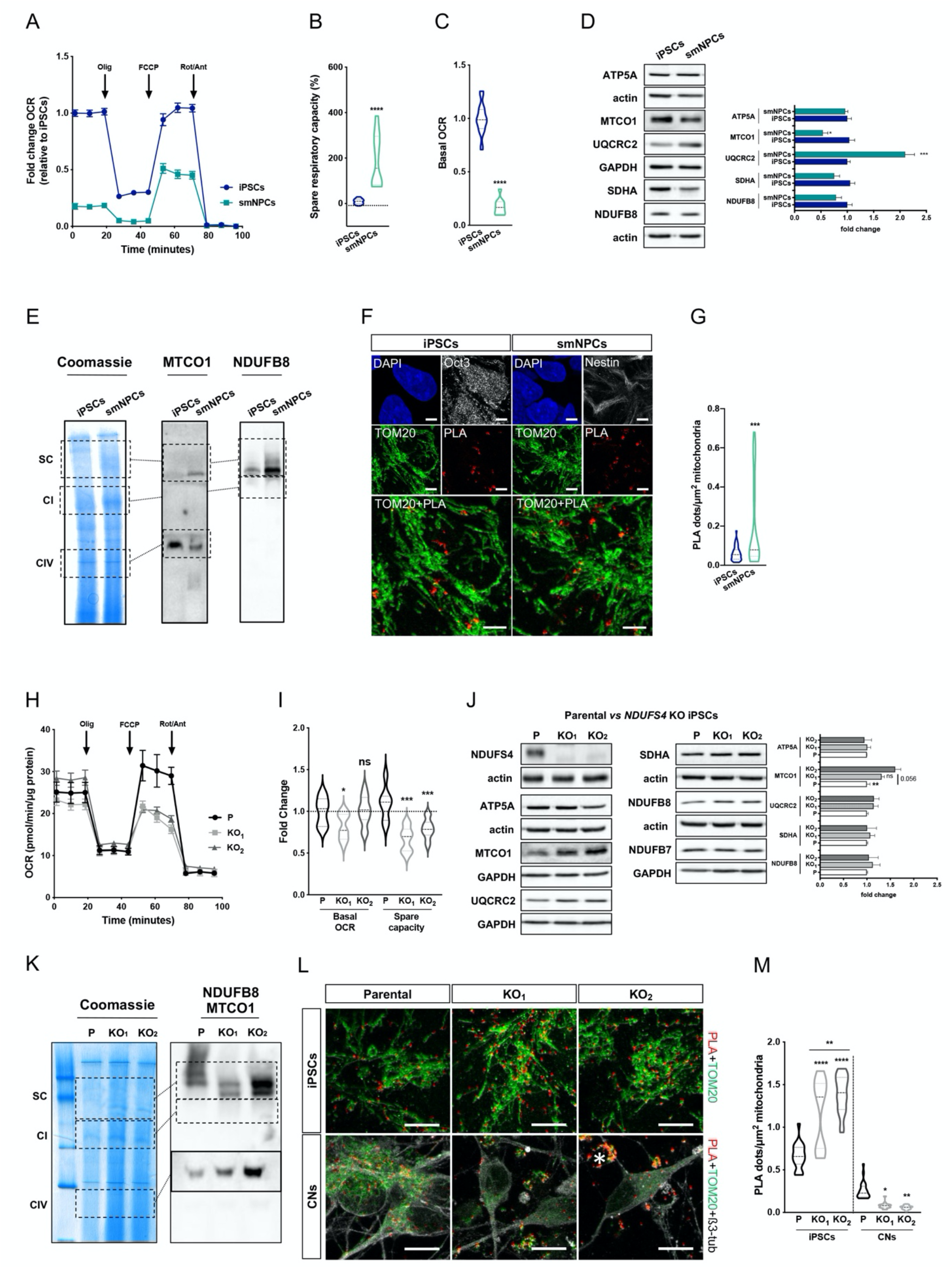
N-respirasome remodeling occurs during iPSC differentiation. (A) OCR measurement of iPSCs and iPSC-derived smNPCs using a conventional Seahorse protocol (n=3). (B) Mitochondrial spare respiratory capacity in human iPSCs and smNPCs. Percentage is relative to their respective basal OCRs (Student’s *t* test, *\*\*\*\*p*<0.0001). (C) Basal OCR in iPSCs and smNPCs (Student’s *t* test, *\*\*\*\*p*<0.0001). (D) Immunoblots using antibodies against OXPHOS system components in iPSCs and smNPCs. Densitometry is relative to iPSCs and reported as mean ± SEM (n=3, Student’s *t* test, \*\**p*<0.01, \**p*<0.05). (E) BN-PAGE and corresponding immunoblots using antibodies against MTCO1 and NDUFB8. (F) Representative pictures of iPSCs and smNPCs stained with DAPI (nucleus, blue), TOM20 (mitochondria, green) and PLA (N-respirasomes, red). Oct3 and nestin staining (both in grey) were used as markers of pluripotency and differentiation, respectively. Scale bar: 5 μm. (G) Quantification of PLA dots normalized to mitochondrial area in iPSCs and smNPCs (n=3; iPSCs = 20 cells; smNPCs = 17 cells; Student’s *t* test, \*\*\**p*<0.001). (H) OCR measurement of wt (P) and two independent clones of *NDUFS4* KO iPSCs (KO1 and KO2). (I) Basal OCR and mitochondrial spare capacity of wt and *NDUFS4* KO iPSCs (fold change, relative to wt; two-way RM ANOVA \*\*\**p*<0.001, \**p*<0.05). (J) Immunoblot analysis of homogenates from parental and *NDUFS4* KO iPSCs using antibodies against ETC subunits (CI, NDUFS4, NDUFB7and NDUFB8; CII, SDHA; CIII, UQCRC2; CIV, MTCO1; CV, ATP5A; actin and GAPDH as loading controls; n=4; Student’s *t* test, \*\**p*<0.01). (K) BN-PAGE and corresponding immunoblots using antibodies against MTCO1 and NDUFB8. (L) Representative confocal images of parental and *NDUFS4* KO iPSCs and immature cortical neurons (CNs). Fluorescent stainings of TOM20 (green) and β3-tubulin (grey) were used as mitochondrial and neuronal markers, respectively. PLA dots are in red. A dying cell is indicated with a white asterisk. (M) Quantification of PLA dots normalized to the mitochondrial area in parental and *NDUFS4* KO iPSCs and immature cortical neurons. Data are represented as a violin plot with median [25^th^ and 75^th^ percentile], one-way ANOVA, n=3, \*\*\*\**p*<0.0001, \*\**p*<0.01, \**p*<0.05.

We wondered whether mitochondrial lesions would alter RSC formation in iPSCs and iPSC-derived cells. To test whether our technique could detect stalled RSC intermediates, we generated *NDUFS4* KO iPSCs using CRISPR/Cas9-gene editing. We obtained two independent lines (indicated as KO_1_ and KO_2_) that carried a homozygous single nucleotide deletion, with a consequent frameshift mutation leading to a premature stop codon (Supplementary Figure S4A-B). Compared to control, NDUFS4 loss led to a mild decrease of basal respiration and a much lower mitochondrial respiratory capacity (Figure 5H-I), although only minor expression changes of other ETC subunits were detected (Figure 5J). When we ran BN-PAGE with freshly prepared samples, we detected superassembled species at lower molecular weight in *NDUFS4* KO cells compared to parental ones (Figure 5K). Using our PLA-based method, we quantified the abundance of N-respirasomes and, in line with the increased amount of superassembled species in the BN-PAGE (Figure 5K), found a significant increase of PLA dots/μm^2^ of mitochondria in *NDUFS4* KO iPSCs compared to parental ones (Figure 5L-M). A possible explanation of this observation is that the incorporation of NDUFS4 in partially superassembled CI occurs at a later stage and is necessary for the stability and optimal functionality of respirasomes as previously shown (Calvaruso et al., 2012; Mimaki et al., 2012; Moreno-Lastres et al., 2012). To further understand the impact of *NDUFS4* KO on N-respirasomes, we differentiated iPSCs into cortical excitatory neurons using transcription factor-mediated forward programming (Peitz et al., 2020). While *NDUFS4* KO did not alter iPSC proliferation (data not shown), it severely undermined the survival of immature cortical neurons, with many cells undergoing degeneration and death (Figure 5L). We performed immunostaining of 6-day-old neurons and found that NDUFS4 KO impaired the formation of N-respirasomes in immature cortical neurons (Figure 5L-M). Thus, although *in-situ* N-respirasome quantification may be considered a suitable proxy marker of mitochondrial functionality, our data in KO iPSCs emphasize that caution is necessary when conclusions are not corroborated by multiple complementary analyses.

### Single cell quantification of N-respirasomes is suitable for imaging-based screens

To determine whether our method can be adapted to imaging-based screens, we performed proof-of-principle experiments using different cultured cellular models. Firstly, we generated murine neural progenitor cells (NPCs) from adult male mice carrying an *Aifm1* allele flanked by attP/attB sites (Wischhof et al., 2018). Upon PhiC31 overexpression and homologous recombination, we genetically deleted *Aifm1 (R200 del)* KI allele in NPCs (Figure 6A). Within a subpopulation of cells overexpressing GFP-tagged PhiC31 and, therefore, lacking *Aifm1*, we could detect an obvious decreased content of N-respirasomes compared to controls (Figure 6B-C). Secondly, we tested whether we could quantify the N-respirasome content in a culture consisting of differentially treated cells. To do so, we transfected HeLa cells with scramble and siRNA against *AIFM1*, the latter inhibiting AIF expression (Figure 6D) and causing OXPHOS impairment and altered OCR (data not shown). We found that siAIF led to a decreased content of N-respirasomes compared to scramble-treated cells (Figure 6E-H). Having confirmed that, we cultured HeLa cells overexpressing mitochondria-targeted GFP and transfected them with scramble siRNA. Then, we mixed these cells with AIF deficient HeLa cells (Figure 6I). Using GFP expression as a cellular marker, we could discriminate the two cell subpopulations and quantify the respective N-respirasome content. As expected, scramble transfected GFP-positive cells (control) had more PLA dots/cell compared to AIF (siAIF) deficient HeLa (Figure 6J-K). This set of data already emphases the potential adaptability of our approach for image-based screens at single cell level. Thirdly, we reasoned that our method could be used to identify novel regulators of N-respirasomes. Thus, we performed tandem mass tags (TMT)-based quantitative proteomics of mitochondria-enriched preparations from hippocampi and cerebella of control and Hq mutant littermates (n=3 per tissue and genotype). Samples were separated by BN-PAGE, prior to excision of high molecular weight bands (~800-1150 kDa) for downstream TMT-based mass spectrometry (MS) analysis (Figure 6L). Compared to controls, 45 and 79 differentially expressed mitochondrial proteins were identified in hippocampal and cerebella samples of Hq mutant mice, respectively (*p* value ≤ 0.1 and ≥ 25% fold change (log2 FC ≥ ± 0.33) thresholds) (Supplementary Figure S5A-D). Of the 33 differentially expressed proteins common in samples from both hippocampus and cerebellum (Figure 6M), 25 proteins are CI and CIII subunits (Supplementary Figure S5D). The remaining 8 proteins were factors associated with RSCs that were excised and analysed from the polyacrylamide gel (Figure 6M). Among them, we found that the CIV component Cox7a2, which is replaced by Cox7a2l/SCAF1 in superassembled structures (Calvo et al., 2020; Cogliati et al., 2016; Garcia-Poyatos et al., 2020), and the MICOS subunit Chchd3 were significantly downregulated in Hq mutant tissues. As suitable candidates for our assay, we transfected HeLa cells with scramble and validated siRNA against *COX7A2L/SCAF1* and *CHCHD3* (Figure 6N). We then performed PLA staining and quantified N-respirasome content. We report that downregulation of COX7A2L/SCAF1 and CHCHD3 influences the amount of N-respirasomes in HeLa cells (Figure 6O-Q). Taken together, our method enables *in-situ* image-based screens of molecular factors involved in RSC biology.

**Figure 6.**
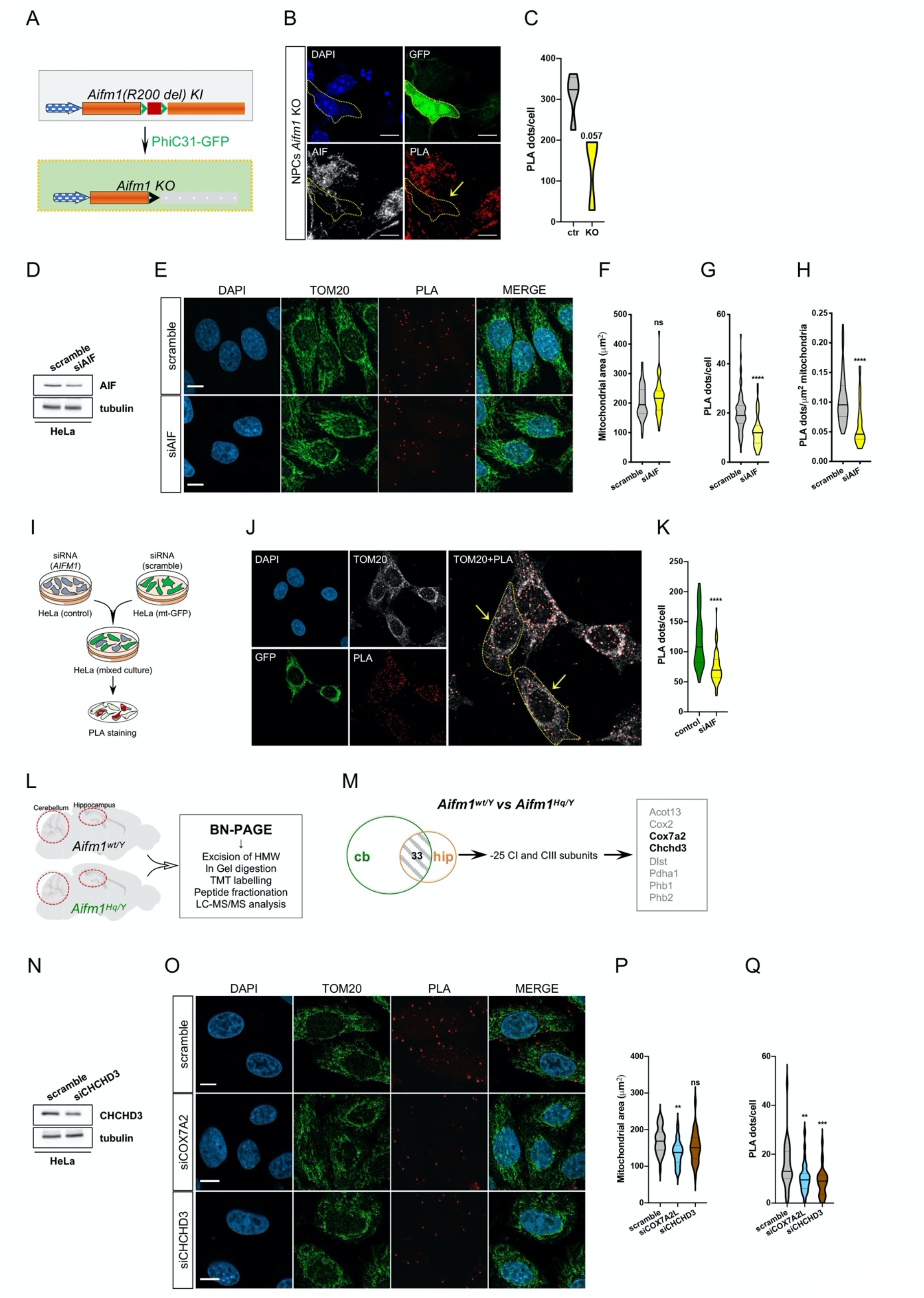
PLA-based imaging analysis of N-respirasomes allows genetic screens in cultured cells. (A) Schematic representation of the *Aifm1(R200 del)* KI allele and its PhiC31-mediated recombination in NPCs. The two att sites (green head arrows) flank exon 5 (red box) in the *Aifm1(R200 del)* KI allele. Upon transfection and overexpression of GFP-tagged PhiC31 recombinase, exon 5 is deleted and AIF expression is impaired. (B) N-respirasomes (PLA, red dots) in AIF KO NPCs generated from *Aifm1(R200 del)* KI mice. Loss of AIF was obtained by transfecting a plasmid encoding a GFP-tagged PhiC31 recombinase. A yellow dot line delineates an AIF KO NPCs (yellow arrow in PLA panel). (C) Quantification of PLA dots/cell in control and AIF KO NPCs. (D) Western blot analysis of HeLa cells transfected with scramble and siRNA against *AIFM1* (siAIF). Immunoblots were developed using antibodies against AIF and tubulin (as a loading control). (E-H) Confocal images and quantification of (F) mitochondrial area (μm^2^), (G) PLA dots/cell and (H) PLA dots/μm^2^ mitochondria of HeLa cells transfected with scramble and siRNA against *AIFM1* (n= 1, scramble= 70 cells, siAIF= 47 cells, Student’s *t* test, \*\*\*\**p*<0.0001). (I) Schematic representation of the experiment. Control HeLa cells were transfected with siRNA against *AIFM1* for 24 hours, while mitochondria-targeted GFP overexpressing HeLa cells (mt-GFP) were transfected with scramble siRNA. Cells were then resuspended, mixed and seeded on the same coverslip. After 24 hours, cells were fixed with PFA and stained for TOM20 and N-respirasomes. (J-K) Representative confocal images and (K) quantification of PLA dots/cell in scramble-treated (control) and *AIFM1* deficient (GFP negative, siAIF) cells, the latter highlighted by yellow dot lines and arrows (n = 1, control = 44 cells, siAIF = 60 cells, Student’s *t* test, \*\*\*\**p*<0.0001). DAPI was used to stain nuclei. (L) Work flow scheme of TMT-based MS analysis. High-molecular weight bands (~800-1150 kDa) from BN-PAGE of wt and Hq mutant hippocampus and cerebellum samples were excised and processed for LC-MS/MS analysis. (M) List of associated factors identified in excised RSCs, differentially expressed in control and Hq mutant cerebella and hippocampi. Of the 33 proteins commonly detected in samples from cerebella and hippocampus, 25 were CI and CIII subunits. Among the other 8 proteins, Cox7a2 and Chchd3 (in bold) were considered for further characterization. (N) Western blot analysis of HeLa cells transfected with scramble and siRNA against *CHCHD3* (siCHCHD3). Immunoblots were developed using anti-CHCHD3 and anti-tubulin antibodies. (O-Q) Representative confocal images and quantification of (P) mitochondrial area (μm^2^) and (Q) PLA dots/cell in HeLa cells transfected with scramble and siRNA against *COX7A2L* and *CHCHD3* (n = 1, scramble = 34 cells, siCOX7A2L = 32 cells, siCHCHD3 = 29 cells; one-way ANOVA \*\*\**p*<0.001, \*\**p*<0.01). Scale bar = 10 μm.

## Discussion

Mitochondrial OXPHOS generates large amounts of ATP and contributes to the maintenance of optimal NAD^+^/NADH ratio in eukaryotic cells. At the crossroad of major metabolic pathways (Spinelli and Haigis, 2018), mitochondria host closely intertwined biochemical reactions that provide carbon units for fatty acid synthesis and gluconeogenesis as well as intermediates for the biogenesis of heme, nucleotide precursors, steroids, amino acids and their derivatives. As a dynamic network of highly interconnected organelle units, mitochondria regulate cell death programs and intracellular calcium homeostasis (Berliocchi et al., 2005; Orrenius et al., 2003), thus integrating endogenous and exogenous cues into spatiotemporal defined signals that drive cellular adaptations and responses in different contexts. Since the maintenance and regulation of mitochondrial bioenergetics critically influence cellular homeostasis, a careful profiling of mitochondrial respiratory integrity serves as a sensitive indicator of the cellular metabolic state. Consistently, a comprehensive overview of mitochondrial paraments is particularly informative for understanding pathophysiological processes as well as for disease diagnosis and prognosis (Connolly et al., 2018; Frazier et al., 2019; Gorman et al., 2016; Koopman et al., 2016; Spinelli and Haigis, 2018).

We herein describe a method for an *in-situ* detection of N-respirasomes in fixed cells as well as in tissues. Our newly established approach takes advantage of the physical distance between CI and CIV subunits when incorporated into superassembled structures. Given the well-characterized properties of the PLA staining, we provide a consistent line of experimental evidence that supports a clear correlation between N-respirasome numbers, composition and organization of the OXPHOS system and functional changes in mitochondrial respiration. In a variety of biologically relevant settings, we report that single-cell quantification of N-respirasomes is technically feasible and represents a useful index for comparative analyses of mitochondrial ETC integrity and OXPHOS capacity. Our PLA-based protocol enables uniform signal patterns in the form of dot-like structures that, as fluorescent or brightfield stainings, are recognizable and quantifiable in either a manual or a semi-automated manner. In HeLa and near-haploid human HAP1 cells, NPCs, iPSC and iPSC-derived cortical neurons, tractable genetic lesions result in a decreased amount of N-respirasomes that correlates with a general impairment of the OXPHOS system and consequent loss of cell fitness. Consistently, our data obtained in mouse-derived tissues further emphasize the sensitivity of our established assay in comparative investigations of mitochondrial defects associated with pathology. In the case of *Aifm1(R200 del)* KI mice, we were able to detect alterations of the OXPHOS system at single-cell resolution that were observed using conventional biochemical methods (i.e., western blot analysis), but not with Seahorse respirometry. As an additional aspect supporting the robustness of our method, we were able to quantify neuron-specific ETC lesions in the cerebellum of Hq mutant mice, despite the extensive microgliosis that blunted other population-based analyses. Moreover, we effectively correlated the abundance of N-respirasomes in iPSC and iPSC-derived neural progenitor cells. Our data imply that undifferentiated iPSCs have less N-respirasomes, whereas partially committed iPSC-derived smNPC exhibit an increased abundance of RSCs, possibly leading to enhanced mitochondrial spare respiratory capacity. As a consequence of *NDUFS4* loss-of-function, a condition often linked with human pathology, altered N-respirasome content is associated with diminished survival of immature excitatory cortical neurons.

Apart from describing a new method with major technical advantages, our study also addresses some unsettled biological questions. First, we report that our *in-situ* PLA-based method clearly reveals mitochondrial defects in tissues from “pre-symptomatic” (i.e., *Aifm1 (R200 del)* KI with no obvious signs of cell death) as well as symptomatic (i.e., *Ndufs4* KO and Hq mutant) mice, whereas conventional Seahorse respirometry detected only marginal differences. By exploiting the sensitivity of our assay, we could discriminate and quantify cell-type specific loss of mitochondrial ETC integrity in degenerating areas of Hq mutant brains. Based on the present and previous line of evidence (Benit et al., 2008; Wischhof et al., 2018), we can conclude with confidence that these two mouse models of AIF deficiency bear equivalent mitochondrial defects, although they develop divergent neurodegenerative profiles. We speculate that reactive gliosis may be a trigger, rather than an associated feature, of neuronal demise in Hq mutant animals. Second, we describe that the abundance of RSCs, and in particular N-respirasome, depends on the metabolic and differentiation states of the cell, with an increased RSCs in those cells that strongly rely on mitochondrial OXPHOS for energy production. In light of these considerations and our experimentally supported conclusions, we propose our PLA-based method as a valuable and complementary approach for single-cell comparative studies of mitochondrial fitness in complex organs, such as the brain. Third, we screen for novel factors that may regulate N-respirasome content. Starting from a proteomic analysis of proteins associated with superassembled species, we found COX7A2 and CHCHD3 among others. COX7A2 is an 83-amino-acid-long CIV subunit that is replaced by COX7A2L/SCAF1 during RSC biogenesis (Calvo et al., 2020; Cogliati et al., 2016; Garcia-Poyatos et al., 2020). Instead, MIC19/CHCHD3 is a mitochondrial intermembrane protein and core component of the mitochondrial contact site and cristae organizing system (MICOS) (Darshi et al., 2011; Stephan et al., 2020). In line with other published evidence (Calvo et al., 2020; Cogliati et al., 2016; Garcia-Poyatos et al., 2020), COX7A2L/SCAF1 downregulation impairs the formation of N-respirasomes in HeLa cells. Regarding MIC19/CHCHD3, our data emphasize that loss of MICOS components, and therefore cristae structures, may alter the formation of superassembled structures, consistently to what recently report in *Drosophila melanogaster* (Murari et al., 2020).

Our method provides robustness and reproducibility when assessing N-respirasomes at single cell level in fixed tissues. We recognize that our PLA protocol is time-consuming, has only a few cost-benefit advantages and may exhibit a reduced detection range when applied to systems that obtain ATP primarily from glycolysis, as in the case of cultured stem cells and tumorigenic cell lines. We are aware that our experiments were performed exclusively with one pair of highly validated antibodies and our PLA-based assessment of N-respirasomes does not cover all possible superassembled species. Our method focuses primarily on RSC I1+III2+IV1 and other eventual, uncharacterized superassembled structures containing NDUFB8 and MTCO1. Due to technical difficulties, we could not provide a direct and mechanistic evidence that would conclusively elucidate the functional relevance of RSC abundance on mitochondrial oxygen consumption and ATP production as recently suggested (Calvo et al., 2020). Although, feasible and marginally described in our work, we did not pursue the quantitative assessment of N-respirasomes in neuronal projections, such as axons and dendrites. As a final consideration, we acknowledge some limitations of our analysis as a proxy measurement of mitochondrial function/dysfunction. In this regard, our technique might not be able to discriminate between fully functional RSCs and NDUFB8- and MTCO1-containing assembly intermediates that accumulate because of stalled OXPHOS biogenesis. Since this aspect represents a potential confounding factor in discerning defects of mitochondrial bioenergetics, we set out to perform experiments using *NDUFS4* KO iPSCs, knowing that NDUFS4 is part of the CI N-module and its incorporation occurs at a later stage and does not undermine the formation of RSC intermediates (Moreno-Lastres et al., 2012). In cells lacking NDUFS4 but still expressing normal levels of other OXPHOS subunits (e.g., i.e., ATP5A, MTCO1, UQCRC2, SDHA, NDUFB8 and NDUFB7), we report that the increased number of N-respirasomes correlates with the enhanced formation of supercomplex assembly intermediates. Although these data may sound surprising, they are consistent with previous evidence elucidating the incorporation of ETC subunits at different stages (Lobo-Jarne et al., 2020; Moreno-Lastres et al., 2012; Protasoni et al., 2020). At least in undifferentiated iPSCs, Seahorse measurements could detect defects in mitochondrial bioenergetics, whereas our study of N-respirasomes would suggest a different scenario. Thus, we call for attention when interpreting PLA-based N-respirasome quantifications that are not corroborated with complementary methods.

Taken together, our findings using *in vitro* cell cultures and preclinical mouse models, of mitochondrial diseases imply that single-cell assessment of N-respirasomes is a quantifiable parameter of mitochondrial respiratory integrity. We envision that this method is suitable for a wide range of biomedical studies and may be adapted for diagnostic screenings of patient-derived materials.

## Materials and methods

### Antibodies and PLA probes

The following primary and secondary antibodies were used in this study: rabbit anti-TOM20 (Santa Cruz Biotechnology, sc-11415); rabbit anti-NUDFB8 (Proteintech, 14794-1-AP); mouse anti-MTCO1 (Abcam, ab14705); anti-calbindin D-28k (Swant, 300); mouse anti-VDAC1/porin (Abcam, ab14734); mouse anti-actin (Sigma, A5316); rabbit anti-AIF (Santa Cruz, sc-13116); mouse anti-total OXPHOS antibody cocktail (Abcam, ab110413); rabbit anti-COX IV (Cell Signaling, 4844); mouse anti-NeuN (Merck Millipore, MAB377); rabbit anti-GAPDH (Santa Cruz, sc-25778); mouse anti-ßIII tubulin (Promega, G7121); mouse anti-NDUFA9 (abcam, ab14713); mouse anti-UQCR2 (abcam, ab14745); rabbit anti-UQCRC2 (abcam, ab127872); goat anti-mouse IgG highly cross-adsorbed secondary Antibody, Alexa Fluor 633 (Invitrogen/ThermoFisher, A-11008); goat anti-rabbit IgG highly cross-adsorbed secondary Antibody, Alexa Fluor 633 (Invitrogen/ThermoFisher, A-21070); goat anti-mouse IgG, IgM (H+L) secondary Antibody, Alexa Fluor 488 (Invitrogen/ThermoFisher, A-10680); goat anti-rabbit IgG, IgM (H+L) secondary Antibody, Alexa Fluor 488 (Invitrogen/ThermoFisher); goat anti-mouse IgG (H+L) highly cross-adsorbed secondary antibody, Alexa Fluor 568 (Invitrogen/ThermoFisher, A-11031); goat anti-rabbit IgG (H+L) cross-adsorbed secondary antibody, Alexa Fluor 568 (Invitrogen/ThermoFisher, A-11011). The following PLA probes were used in this work: Duolink In Situ PLA Probe Anti-Mouse Minus (SIGMA, DUO9004); Duolink In Situ PLA Probe Anti-Mouse Plus (SIGMA, DUO92001); Duolink In Situ PLA Probe Anti-Rabbit Minus (SIGMA, DUO92005); Duolink In Situ PLA Probe Anti-Rabbit Plus (SIGMA, DUO92002); Duolink In Situ Detection Reagents Red (SIGMA, DUO92008); Duolink In Situ Detection Reagents Brightfield (SIGMA, DUO92012).

### Cell culture

Near-haploid HAP1 cells (Horizon Genomic’s GmbH, Vienna, Austria) were grown in Iscove’s modified Dulbecco medium (IMDM) supplemented with 10% fetal bovine serum and 1% penicillin/streptomycin (100 U/mL penicillin; 100 μg/mL streptomycin). Parental and mitochondrial mutant HAP1 cells were maintained and used as previously described (Gioran et al., 2019). Drug treatments with oligomycin, FCCP, rotenone/antimycin A in HAP1 cells were carried out for 30 min following the concentrations used for Seahorse experiments (Connolly et al., 2018). iPSCs were cultured under feeder-free conditions in StemMACs iPS-Brew (Miltenyi Biotec) on vitronectin-coated 6-well plates with daily medium changes. For OCR measurements (see below), iPSCs were split with EDTA and 5×10^4^ cells per well were seeded on Seahorse XF24 cell culture microplates (Agilent) 24 h before the experiment. For PLA-experiments, 1×10^5^ cells per well were seeded on vitronectin-coated coverslips in 12-well plates, and fixed with 4% PFA on the next day. iPSCs-derived small molecule neural precursor cells (smNPCs) were cultured on Matrigel-coated 6-well plates and kept in N2B27 medium consisting of DMEM-F12 (Invitrogen)/Neurobasal (Invitrogen) with N2 and B27 (without vitamin A) supplement, 1% glutamine, 0.5 μM Purmorphamine (Miltenyi Biotec), 3 μM CHIR 99021 (Miltenyi Biotec) and 150 μM LAAP (Merck). Medium was replaced every other day. For OCR measurements, smNPCs were split with accutase and 2×10^5^ cells per well were seeded on Seahorse cell culture microplates (Agilent) one day before the, experiment. For PLA experiments, 3×10^5^ cells per well were seeded on Matrigel-coated coverslips in 12-well plates and fixed with 4% PFA 48 to 72 h after seeding. smNPCs were spontaneously differentiated by withdrawal of supplements from the N2B27 medium. After three weeks, cultures were mitotically inactivated by treatment with 5 μM AraC, resulting in a predominant neuronal population with less than 1% glia cells (data not shown). Cells were differentiated for a total of 4 to 5 weeks with medium changes every other day. For PLA experiments, 1×10^5^ smNPCs per well were seeded on Matrigel-coated coverslips in 12-well plates, switched to differentiation medium the next day, and fixed with 4 % PFA after 4 to 5 weeks of differentiation.

HeLa cells were grown in DMEM supplemented with 10% fetal bovine serum and 1% penicillin/streptomycin (100 U/ml penicillin; 100 μg/ml streptomycin). For lipid-mediated siRNA transfection, cells were trypsinized and seeded onto glass coverslips in 24-well plates (3 × 10^4^ cells/well) together with transfection complexes (Lipofectamine 3000 and siRNAs at a final concentration of 100 nM) diluted in Opti-MEM. After 48 h, cells were fixed in 4% PFA and used for PLA experiments. The following siRNAs were used: negative control AM4611, siAIF ID117234, siCHCHD3 ID147671, siCOX7A2L ID64381, siNDUFS3 ID224103 (Ambion, Thermo Fisher).

### Generation of NDUFS4 KO iPSCs

To generate *NDUFS4* KO iPSCs, we used human iPS cell line C-14m-s11-NGN2, which carries a doxycycline-inducible NGN2 transgene that enables rapid forward programming into excitatory neurons (Rhee et al., 2019). Cells were nucleofected with a CRISPR-Cas9 ribonucleoprotein (RNP) complex containing Alt-R HiFi Cas9 Nuclease (IDT) and a tracrRNA:crRNA duplex (crRNA sequence: GGTCGTTGAGGACTTCCACA) using program CM150 of the Amaxa 4D Nucleofector (Lonza). Briefly, the transfection solution (16.4 μl P3 solution, 3.6 μl supplement 1, 0.5 μl electroporation enhancer and 2.5 μl RNP mix) consisted of equal volumes of Cas9 protein (from 61 μM stock) and an annealed tracrRNA:crRNA duplex solution (all reagents were purchased from IDT, USA). Nucleofection was performed with 20 μl of the transfection solution. From the resulting genome-edited polyclonal cell line, single cell-derived sub-clones were isolated. Frameshift-inducing mutations were identified by Sanger sequencing. Single clones were selected if they showed a Knockout Score of at least 99% according to the ICE CRISPR Analysis Tool (www.synthego.com). Sequence-validated clones were further expanded and quality-controlled by virtual karyotyping using SNP arrays and regular mycoplasma testing. Candidate C-14m-s11-NGN2 clones were then differentiated into excitatory neurons following a previously published protocol (Peitz et al., 2020).

### Human postmortem tissues

Human brain sections were obtained from The Netherlands Brain Bank (NBB), Netherlands Institute for Neuroscience, Amsterdam. All Material has been collected from donors for or from whom a written informed consent for a brain autopsy and the use of the material and clinical information for research purposes had been obtained by the NBB.

### Imaging analysis and PLA quantification

PLA fluorescence-stained cells were imaged using the next generation confocal microscopes with Airyscan Zeiss LSM800 or LSM900 with a 63× oil immersion objective. All the PLA measurements were obtained using z-stacks of 5-8 images (5 images for cells in vitro, 8 images for cells in tissues) with a thickness of 0.5 μm between each focal plane. Images were deconvolved in ZEN blue (Zeiss) and maximum projections were obtained using Fiji/ImageJ, which was used also for the semi-automatic quantitative assessment of PLA-dots. (A) For cultured cells, images were taken with ROIs containing 6-10 cells. Single cells were cropped and, based on the TOM20 staining, a mitochondrial mask (gaussian blur filter, subtract background, auto threshold “method=default”) was created to measure the mitochondrial area of the cell. PLA-dots were counted in the mask using the Fiji/ImageJ option “find maxima”, excluding the possible off-target signals outside the mitochondrial network. PLA-dots were normalized based on mitochondrial area. (B) For fixed tissues, images were taken with ROIs containing 1-5 cells. Single cells were cropped and, based on cell (NeuN for CA1 neurons, calbindin for cerebellar Purkinje cells) and mitochondrial markers (TOM20), cell body and mitochondrial area were measured by applying a mask, as described above. PLA-dots were counted, as described for the cell cultures, and normalized based on the area of the soma or mitochondria. PLA Brightfield images were obtained using an episcope apotome microscope with a 20× air objective.

### Oxygen consumption rate measurements

Assessment of the oxygen consumption rate (OCR) in mouse brain tissue was performed as previously described (Fried et al., 2014), with slight modifications. Briefly, acute brain sections were prepared from transgenic and control mouse littermates. Animals were briefly anesthetized with Isoflurane and then sacrificed by cervical dislocation. Brains were rapidly removed and placed into pre-warmed artificial cerebrospinal fluid (aCSF: 120 mM NaCl, 3.5 mM KCl, 1.3 mM CaCl_2_, 1 mM MgCl_2_, 0.4 mM KH_2_PO_4_, and, 5 mM HEPES with freshly added 10 mM glucose). Then, brains were sliced into 1 mm thick sections using a rodent brain slicer (Zivic Instruments, Pittsburgh, USA). Sections were kept in pre-warmed aCSF throughout the entire experimental procedure. A 5 ml pipette tip was then used to obtain consistent tissue punches from the cortex and cerebellum. Samples were loaded onto an XF Islet Capture Microplate (101122-100, Agilent Seahorse) and secured in the center of the well with mesh capture screens. Pre-warmed aCSF-medium (700 μl aCSF, supplemented with 4 mg/ml BSA) was immediately pipetted into each well. The microplate was then incubated at 37 ℃ for 1 h in a CO_2_-free incubator to allow for temperature and pH equilibration. The OCR was then measured with an XF24 Extracellular Flux Analyzer via the XF Cell Mito Stress Kit (Agilent Seahorse). Following 5 measurements of basal OCR, the ATP synthase inhibitor oligomycin (2 μM) as well as the mitochondrial uncoupler carbonyl cyanide 4-(trifluoromethoxy)-phenylhydrazone (FCCP, 2 μM) and the complex I/II inhibitor mix rotenone/antimycin A (0.5 μM) were successively added.

Cultured cells were seeded on cell culture microplates in culture medium 24 h before the assay. On the day of the experiment, growth media were replaced with Seahorse XF base medium supplemented with 1 mM pyruvate, 10 mM glucose, and 2 mM glutamine. Before the measurement, cells were equilibrated for 60 min in a CO_2_-free incubator at 37 ºC. Following three baseline measurements, oligomycin, FCCP and rotenone/antimycin A were successively added. The following concentrations were used: HAP1 cells: 1 μM oligomycin, 0.5 μM FCCP, 0.5μM rotenone/antimycin A; iPSCs: 0.5 μM oligomycin, 1 μM FCCP, 0.5 μM rotenone/antimycin A; smNPCs: 1 μM oligomycin, 1 μM FCCP, 0.5 μM rotenone/antimycin A. At the end of the OCR assessment, cells were collected and lysed in RIPA buffer (SIGMA), supplemented with protease and phosphatase inhibitors (Roche). Protein concentrations were determined via Bradford assay. OCR values were then normalized to the respective protein contents.

### PLA fluorescence

Cells were washed once in PBS, fixed with 4% PFA for 5 min at 4 °C, rinsed 3× 5 min with PBS. Fixed cells were permeabilized with 0.25% Triton in PBS for 1h at RT, rinsed 2× 5 min with PBS and treated with H_2_O_2_ (3% in dmH_2_O, 50 μl/coverslip) for 30 min at RT. Cells seeded on glass coverslips were washed 2× 5 min with PBS and incubated with 50 μl Blocking Solution for 1 h at 37 °C. Samples were incubated with primary antibodies (MTCO1 1:200 and NDUFB8 1:100, diluted in antibody diluent, 50 μl/coverslip) in a humidity chamber overnight at 4 °C, washed 2× 5 min with pre-warmed PLA wash buffer A and finally incubated, with PLA Probes 5× (in antibody diluent, 50 μl/coverslip) for 1 h at 37 °C. After 2× 5 min washing with Wash buffer A, ligation reaction was carried out by incubating with the Ligation-Ligase solution (in mqH_2_O, 50 μl/sample) for 1 h at 37 °C. Amplification reaction was performed after 2× 2 min with PLA Wash buffer A, by incubating with the Amplification-Polymerase solution (in mqH_2_O, 50 μl/sample) for 2 h at 37 °C.

Brain slices were washed once in PBS while agitating, incubated with citrate buffer (pH 6.0) for 30 min at 90 °C, then for another 30 min at RT. Tissues were washed 2× 5 min in PBS, permeabilized in a solution containing 0.5% Triton X-100 (in PBS) and 3% H_2_O_2_ (in mqH_2_O, 1 ml/well) for 30 min at RT, washed 2× 5 min with PBS and incubated with Blocking Solution (50 μl/section) for 1 h at 37 °C. Samples were incubated with primary antibodies as described above for 48h at 4 °C, washed 2× 5 min with PLA Wash buffer A while agitating, then incubated with PLA Probes 5× (in antibody diluent, 50 μl/section) for 24 h at 4 °C. After 2× 5 min washing with Wash buffer A, ligation and amplification reactions and development were performed as described above. Cells and brain sections were washed 2× 5 min with PLA Wash buffer B, incubated overnight at 4 °C with the primary antibodies for mitochondria (i.e, TOM20) and cell identity (i.e., NeuN or calbindin) in blocking solution, washed 2× 5 min with PBS 1× and exposed to secondary antibodies for 2 h at RT in the dark. After 2× 5 min washing with 1× PBS, cells and brain sections were incubated with DAPI (1:10.000 in PBS) for 10 min and 30 min at RT in the dark, respectively. Cells-seeded coverslips and brain sections were washed 3× 5 min with PBS and mounted on glass slides using Dako fluorescence Mounting medium. Samples were kept in the dark at 4 °C and imaging analysis were run within 48 h.

### PLA Brightfield

(A) Brain slices were washed once in PBS while agitating, incubated with citrate buffer (pH 6.0) for 30 min at 90 °C, then for another 30 min at RT. Tissues were washed 2× 5 min in PBS, permeabilized in a solution containing 0.5% Triton X-100 (in PBS) for 1 h at RT. Tissue sections were washed 5 min with PBS, permeabilized with 3% H_2_O_2_ (in mqH_2_O, 1 ml/well) for 30 min at RT, washed 2× 5 min with PBS and incubated with Blocking Solution (60 μl) for 1 h at 37 °C. Samples were incubated with primary antibodies as described above for 48 h at 4 °C, washed 2× 5 min with PLA Wash buffer A while agitating, then incubated with PLA Probes 5× (in antibody diluent, 60 μl/section) for 24 h at 4 °C. After 2× 5 min washing with Wash buffer A, ligation reaction was carried out by incubating with the Ligation-Ligase solution (in mqH_2_o, 60 μl/sample) for 1 h at 37 °C. Amplification reaction was performed after 2× 2 min with PLA Wash buffer A, by incubating with the Amplification-Polymerase solution (in mqH_2_o, 60 μl/sample) for 2 h at 37 °C. After 2× 5 min washing steps with PLA Wash buffer A, samples were incubated with the 5× Detection Brightfield Solution (in mqH_2_o, 60 μl/ section) for 1 h at RT. After 2× 2 min washing with mqH_2_O, brain sections were mounted with 0.7% gelatin. The day after, they underwent dehydration procedure consisting of 1× 5 min in 70% ethanol, 1× 5 min in 96% ethanol, 2× 2 min in 100% ethanol, 8 min Xylene and final mounting in DPX medium.

### Sample preparation, LC-MS/MS measurements and database searching

High molecular weight bands (~800-1150 kDa) in blue native gels of hippocampal and cerebellar samples (n=3) from wt and Hq mutant mice, were excised, destained, reduced and alkylated prior to overnight in gel digestion with sufficient Trypsin (20 ng/μl, in 50 mM ammonium bicarbonate) to completely cover the gel pieces. Peptides were extracted from the gel slices by adding 100 μl of extraction buffer (1:2 (vol/vol) of 5% formic acid/ acetonitrile) and incubation at 37 °C with shaking for 15 min. Extracted peptides were dried in a vacuum centrifuge and re-suspended in 15 μl of 50 mM HEPES (pH 8.5). Peptides were processed in 2 batches and labelled with 25 μl of diluted (14.75 mM) 128N, 128C, 129N and 129C, 130N, 130C TMT 10plex reagents, respectively, for 1h at RT. TMT signal was quenched by addition of 2 μl of 5% hydroxylamine to the reaction, vortexing for 20 s and incubating for 15 min at 25 °C with shaking (1000 rpm). TMT-labelled peptides were acidified with 45% (vol/vol) of 10% FA in 10% ACN, prior to combining samples at equal amounts for drying in a vacuum centrifuge. Dried peptides were re-suspended in 300 μl of 0.1% TFA for subsequent high pH reverse phase fractionation (Pierce kit). Then, 6 peptide fractions (10, 15, 20, 25, 50, and 80%) ACN were collected, concentrated and re-suspended in 20 μl of 5% FA for LC-MS analysis.

Tryptic peptides were analyzed on a Dionex Ultimate 3000 RSLC nanosystem coupled to an Orbitrap Exploris 480 MS. Peptides were injected at starting conditions of 95% eluent A (0.1% FA in water) and 5% eluent B (0.1% FA in 80% ACN). Peptides were loaded onto a trap column trapping cartridge (Acclaim PepMap C18 100Å, 5 mm × 300 μm i.d., #160454, Thermo Scientific) and separated by reversed-phase chromatography on a 50 cm μPAC C18 column (PharmaFluidics) using an 85 min linear increasing gradient from 8% to 25% of eluent B followed by a 28 min linear increase to 50% eluent B. The mass spectrometer was operated in data dependent and positive ion mode with MS1 spectra recorded at a resolution of 120k with an automatic gain control (AGC) target value of 300% (3×10^6^) ions, maxIT set to Auto and an intensity threshold of 1×10^4^, using a mass scan range of 350−1550. Precursor ions for MS/MS were selected using a top speed method with a cycle time of 2 ms and normalized collision energy (NCE) of 36% (High-energy Collision Dissociation (HCD)), to activate both the reporter and parent ions for fragmentation. MS2 spectra were acquired at 45k resolution using an AGC target value of 200% (2×10^5^), and maxIT set to 86 ms. Dynamic exclusion was enabled and set at 45 s. Isolation width was set at 0.7 m/z and the fixed first mass to 110 m/z to ensure reporter ions were detected. Peptide match was set to off, and isotope exclusion was on. Charge-state exclusion rejected ions that had unassigned charge states, were singly charged or had a charge state above 5. Full MS data were acquired in the profile mode with fragment spectra recorded in the centroid mode.

Raw data files were processed with Proteome Discoverer™ software (v2.4.0.305, Thermo Scientific) using SEQUEST® HT search engine against the Swiss-Prot® Mus musculus database (v2020-01-15). Peptides were identified by specifying trypsin as the protease, with up to 2 missed cleavage sites allowed. Precursor mass tolerance was set to 10 ppm, and fragment mass tolerance to 0.02 Da MS2. Static modifications were set as carbamidomethylated cysteine and TMT6plex (229.163 Da; N-terminal, K), while dynamic modifications included methionine, oxidation and N-terminal protein acetylation, for all searches. Resulting peptide hits were filtered for maximum 1% FDR using the Percolator algorithm. The TMT10plex quantification method within Proteome Discoverer software was used to calculate the reporter ratios with mass tolerance ±10 ppm and applying isotopic correction factors. Only peptide spectra containing all reporter ions were designated as “quantifiable spectra”.

### Statistics

Data were analysed using Graph Pad Prism and are presented as mean ± SEM. The number of biological replicates is indicated in the figure legends. Statistical analysis of normal distributed data was done with two-tailed Students’s *t* test, one- or two-way ANOVA with repeated measures where appropriate. Non-normal distributed data was statistically compared with Mann-Whitney *U*-Test. Statistical significance was defined as *\*p<*0.05, not significant was indicated as ns.

### Transgenic mice and animal work approval

Aifm1 (R200 del) knockin mice were generated as previously described (Wischhof et al., 2018). B6.129S4-*Ndufs4^tm1.1Rpa^*/J (Jax Stock No 027058 and B6CBACa *A^w-J^*/*A*-*Aifm1^Hq^*/J (Jax Stock No 000501) were obtained from The Jackson Laboratory (Bar harbor, Maine, USA). All mice were housed under a 12/12 h light/dark cycle and fed *ad libitum* with regular Chow diet. All experiments were performed according to the DZNE internal guidelines and approved by the State Agency for Nature, Environment and Consumer Protection in North Rhine Westphalia (LANUV, NRW; application numbers: 84-02.04.2014.A521 and 81-02.04.2020.A110).

## Acknowledgements

We wish to thank our DZNE colleagues at LMF; Ms. Christiane Bartling-Kirsch for her assistance; Prof. Donato Di Monte and Prof. Werner Koopman for their critical comments; The Netherland Brain Bank (NBB) for the human *postmortem* tissues kindly provided for this work. This research was supported by the DZNE institutional budget, the CoEN (Carbon-Model, 3018) initiative and the Helmholtz cross-program topic “Aging and Metabolic Programming (AMPro)”. Dr. Fabio Bertan and Ms. Anaïs Marsal-Cots received funding from the European Union’s Horizon 2020 research and innovation programme under the Marie Skłodowska-Curie grant agreement No 676144 (Synaptic Dysfunction in Alzheimer Disease, SyDAD). LW, PN and DB are members of the DFG Cluster of Excellence ImmunoSensation funded by the Deutsche Forschungsgemeinschaft (DFG, German Research Foundation) under Germany’s Excellence Strategy – EXC2151 – 390873048. JJ, JHMP and DB are members of the Mitochondrial Dysfunction in Parkinson’s Consortium (PD-MitoQUANT). PD-MitoQUANT has received funding from the Innovative Medicines Initiative 2 Joint Undertaking under grant agreement No. 821522. This Joint Undertaking receives support from the European Union’s Horizon 2020 research and innovation programme and EFPIA.

## Author contribution

Conceptualization: FB, LW and DB; Validation, Formal Analysis and Investigation: FB, LW, ES, MG, JJ, AMC, AP, MS, MP; Writing: FB, DB; Visualization FB, LW and DB; Supervision, FB and DB; Project Administration, DB; Funding Acquisition, MP, JHMP, PN and DB.

## Conflict of interest

The authors declare no conflict of interest.

---

**Supplementary Figure S1 (relative to figure 3).**
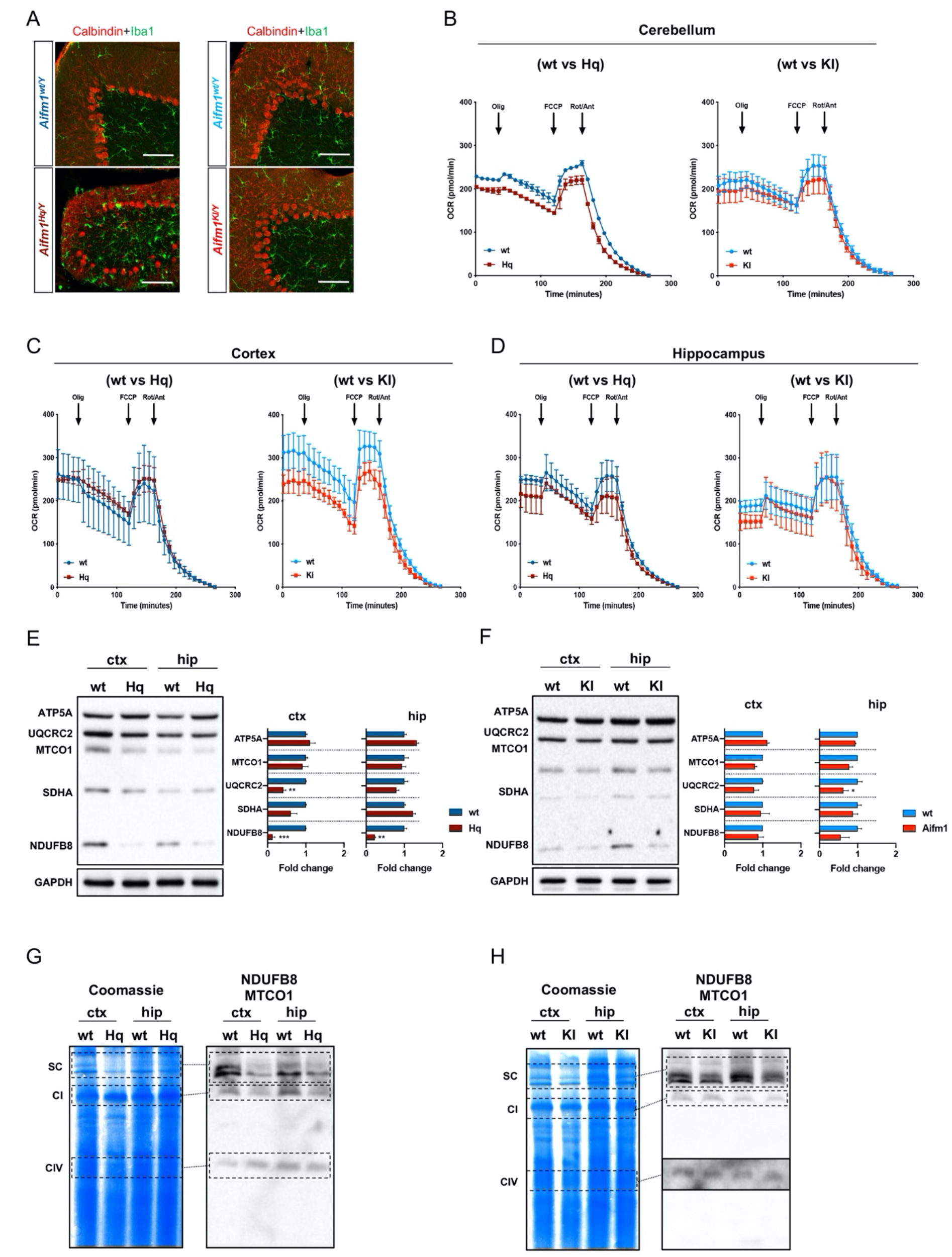
(A) Low-magnification confocal images of cerebella stained with calbindin (Purkinje, red) and Iba1 (microglia, green) in control (*Aifm1^wt/Y^*), Harlequin (*Aifm1^Hq/Y^*) and *Aifm1 (R200 del*) knockin (*Aifm1^KI/Y^*) mice. Scale bar: 50 μm. (B-D) OCR measurements in *ex vivo* (B) cerebellar, (C) cortical and (D) hippocampal sections from 6-month-old mice (wt, n=2; Hq, n=2; wt, n=4; KI, n=4). (E-F) Immunoblot analyses using brain samples from (E) Hq mutant and wt littermates and (F) *Aifm1 (R200 del)* KI and wt littermates (for all genotypes, n=3). Data are represented as fold change (relative to wt), mean ± SEM (\*\*\**p*<0.001, \*\**p*<0.01, \**p*<0.05). (G) BN-PAGE and immunoblots using antibodies against NDUFB8 (CI) and MTCO1 (CIV). Samples are cortices and hippocampi from wt and Hq mice (wt, n=3; Hq, n=3). (H) BN-PAGE and composite image of immunoblots (NDUFB8, MTCO1) using freshly frozen samples (ctx: cortex; hip: hippocampus) from wt and *Aifm1 (R200 del)* KI mice.

**Supplementary Figure S2 (relative to Figure 3).**
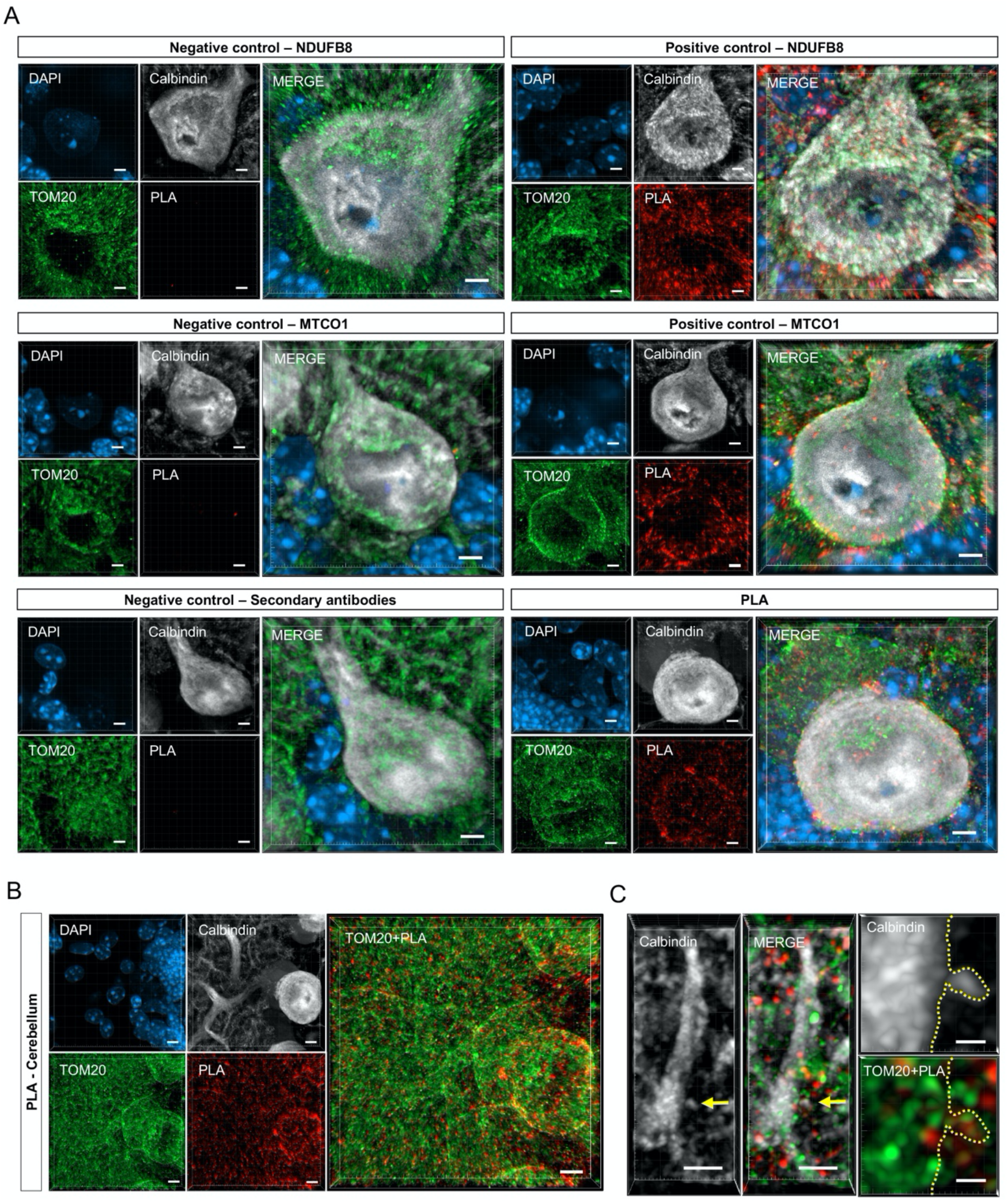
(A) Single-cell validation of PLA method in wt cerebellar brain sections stained with DAPI (nucleus, blue), calbindin (Purkinje cells, gray), TOM20 (mitochondria, green) and PLA (N-respirasomes, red). Negative and positive controls are for antibodies against NDUFB8 and MTCO1 and/or corresponding IgGs. Scale bar: 5 μm. (B) Representative pictures of cerebellar Purkinje cells from 6-month-old control tissue stained with DAPI (nucleus), calbindin (Purkinje neurons), TOM20 (mitochondria) and PLA (N-respirasomes). Single channel scale bar: 20 μm. (C) High-resolution image of stained dendrites and dendritic spines of a Purkinje neuron. Dendrite scale bar: 2 μm; dendritic spine scale bar: 0.5 μm.

**Supplementary Figure S3 (relative to Figure 4).**
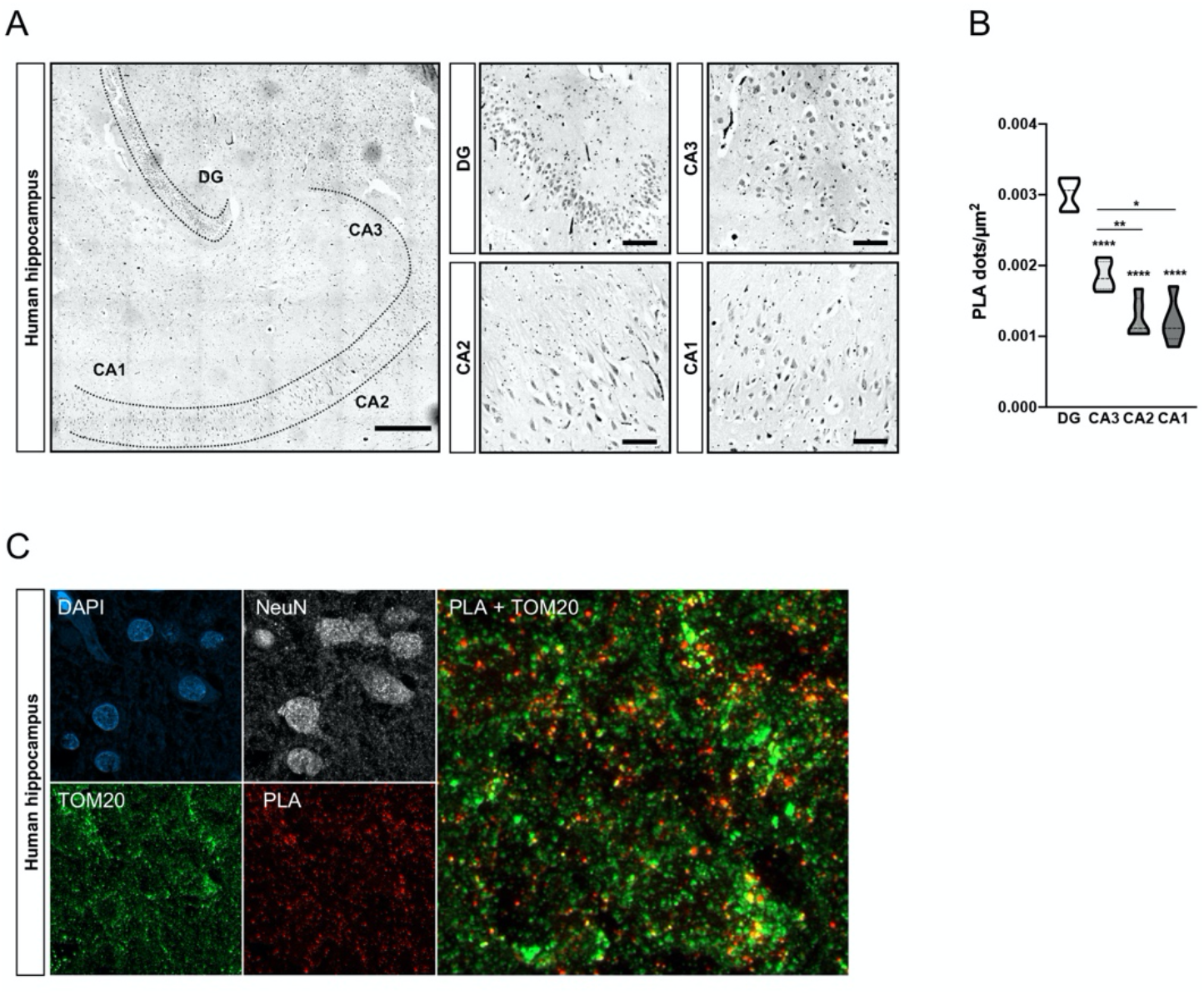
(A) Brightfield PLA of human hippocampus (left panel) and higher magnification images of DG, CA3, CA2 and CA1 (right panels). (B) Quantification of PLA dots/μm^2^ in the indicated areas of the hippocampus (n=1). (C) Fluorescent staining of human hippocampus using NeuN (as a neuronal marker; grey), TOM20 (as a mitochondrial marker; green) and PLA (red).

**Supplementary Figure S4 (relative to Figure 5).**
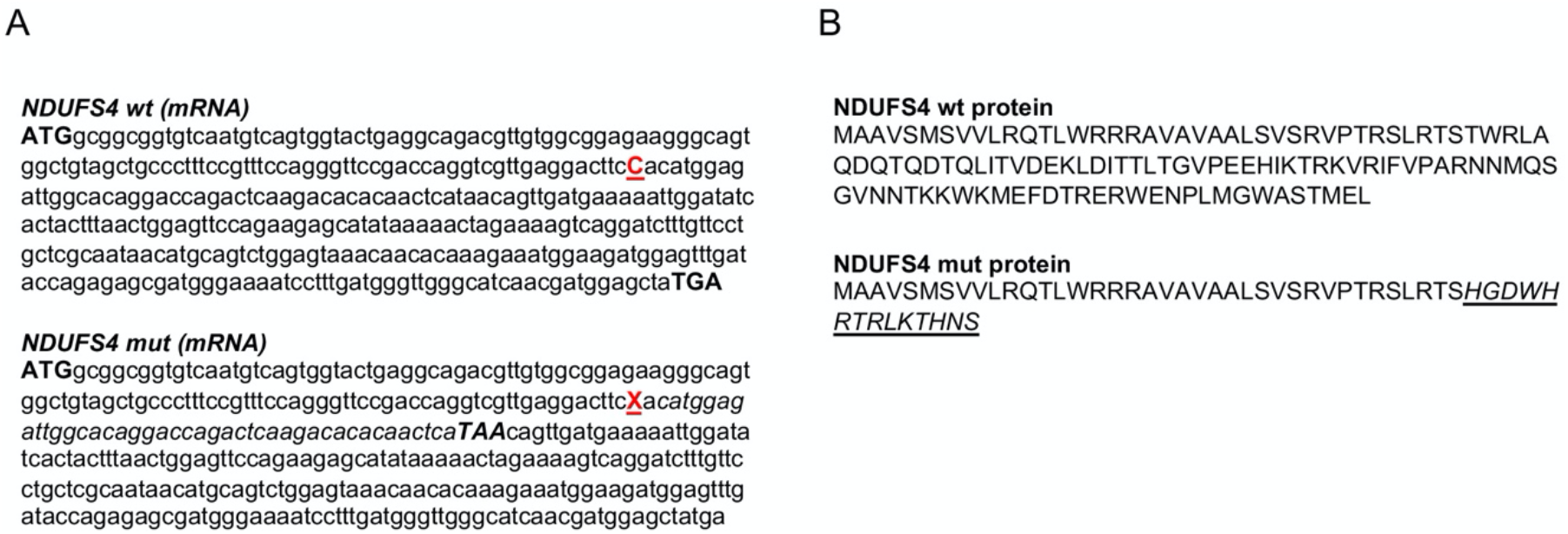
(A) cDNAs from mutated CRISPR/Cas9-gene edited *NDUFS4* alleles were sequenced and aligned with wild type *NDUFS4* transcripts (NCBI reference: XM_017009491.1). A single nucleotide deletion (red C in the *NDUFS4* wt; deletion depicted with a red X in *NDUFS4* mut) led to a premature stop codon (capital italic letters in *NDUFS4* mut). (B) Annotated wild type (top) and mutated NDUFS4 protein sequence (bottom) upon gene editing. The frameshift mutation resulted in a truncated protein with 14 predicted mutated amino acids (capital italic and underlined letters).

**Supplementary Figure S5 (relative to Figure 6).**
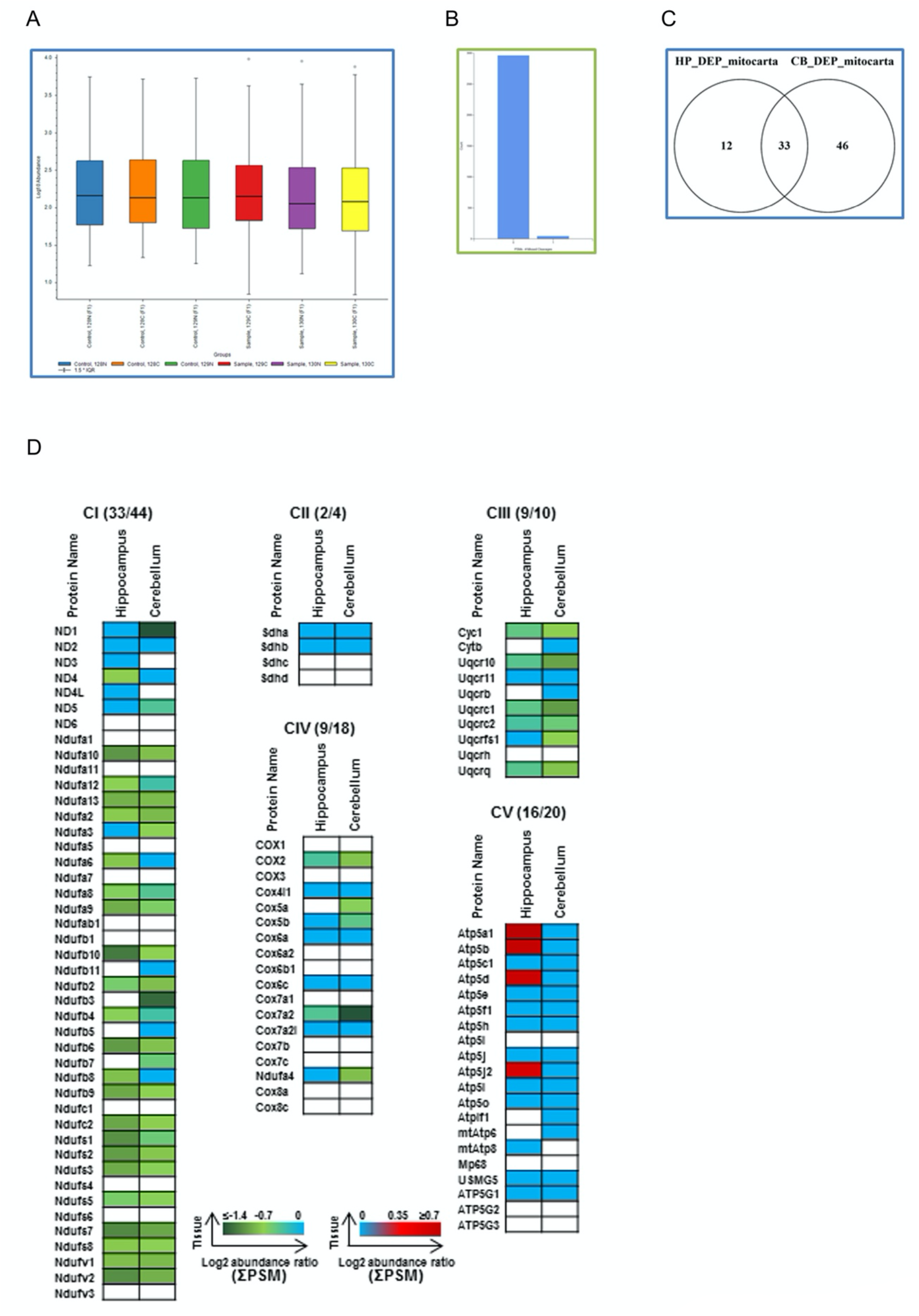
(A) Normalized TMT signal shows comparable total signal levels across the 6 TMT channels (128N, 128C, 129N, 129C, 130N and 130C). (B) Bar chart of missed cleavages after trypsin digestion indicates that the majority of peptides were completely digested by the enzyme and very few had 1 missed cleavage site. (C) Venn diagram summary of differentially expressed mitochondrial proteins (based on Mouse MitoCarta v2.0). In hippocampus and cerebellum samples from Hq mutant mice, 33 proteins were consistently identified. (D) Identified ETC subunits are depicted. Protein changes in Hq mutant mouse tissues are shown in green (downregulated), blue (no change) and red (upregulated). Unidentified proteins are shown in white.

## Notes

### Competing Interest Statement

The authors have declared no competing interest.

